# Evolution of regulatory networks controlling plasticity in gene expression between *Saccharomyces cerevisiae and Saccharomyces paradoxus*

**DOI:** 10.64898/2026.05.18.725926

**Authors:** Anna C. Redhuis, Patricia J. Wittkopp

## Abstract

Organisms cope with environmental changes by modifying gene expression. To understand how regulatory networks controlling expression plasticity evolve, we analyzed RNAseq data from *Saccharomyces cerevisiae*, *Saccharomyces paradoxus*, and their F_1_ hybrids at multiple timepoints after transferring cells from standard laboratory conditions to five environments (low phosphorus, low nitrogen, hydroxyurea shock, heat stress, and cold stress) and during the diauxic shift. In each of the six datasets, we identified genes that changed expression following the transition to the new environment and used hierarchical clustering to identify genes that increased or decreased in expression. We then compared these classifications between orthologs to identify genes with divergent plasticity. For some genes, plasticity was more extreme in one species than the other, and for others, expression of orthologs changed in opposite directions when acclimating to the same environment. Most cases of plasticity divergence were seen only in one environment and were attributable primarily to *trans*-regulatory divergence. Using environment-specific regulatory networks inferred from data in Yeastract, we found that divergent plasticity of environment-specific transcription factors generally did not predict divergent plasticity of their target genes. We also found that, as a group, genes with conserved plasticity tended to have more regulatory interactions than genes with divergent plasticity. Interesting patterns of expression divergence were also observed for five transcription factors in the pleiotropic drug resistance network and their target genes that might contribute to phenotypic divergence. Together, these findings show how environment-specific *trans*-regulatory divergence and combinatorial gene regulation shape the evolution of expression plasticity.

## Introduction

Organisms live in constantly changing environments and have evolved to cope with these environmental changes through phenotypic plasticity, which is the ability of a single genotype to produce alternate phenotypes under different environmental conditions. Plasticity is controlled at the molecular level by changes in the regulation of gene expression. This regulation depends upon interactions between *cis*-acting DNA sequences (e.g., enhancers, promoters) and *trans*-regulatory RNAs and proteins (e.g., transcription factors, signaling proteins). Evolutionary divergence in either *cis*-regulatory sequences or sequences encoding *trans*-regulatory factors can alter the regulation of gene expression. Changes in expression plasticity between populations and species can be important sources of ecological diversification (Hodgins-Davis and Townsend 2009; Lasky et al. 2014; Ghalambor et al. 2015; Makinen et al. 2018; Ballinger et al. 2023). Studies examining the evolution of expression plasticity often focus on responses to a single environmental shift, but organisms experience many different environmental shifts during their lifetimes. Understanding how the same genetic divergence affects expression plasticity across a range of environments is thus essential to understanding how gene regulatory systems evolve.

Expression plasticity begins when a cell detects a shift in the environment, for example via depletion of a nutrient or a change in temperature. This altered cellular state then triggers a change in gene expression. But expression plasticity is not simply a gene being “on” or “off” in different environments. Rather, expression plasticity is how much and how quickly expression is activated or repressed in response to a change in environment (Hager et al. 2009; Rivera et al. 2021; Wagh et al. 2023). Divergence in expression plasticity is therefore best thought of as shifts in these expression dynamics between species (Koster et al. 2015). Detecting such changes requires measuring gene expression in the environment(s) that reveal the divergence in plasticity, whereas detecting genetic changes that affect a gene’s expression level generally can be done in a single environment (Figure 1A). Measuring gene expression in multiple environments begins to disentangle changes in expression level from expression plasticity (Figure 1B,C), whereas measuring expression at multiple time points following multiple environmental changes provides insight into the dynamics of the plastic expression response (Figure 1D,E).

**Figure 1.**
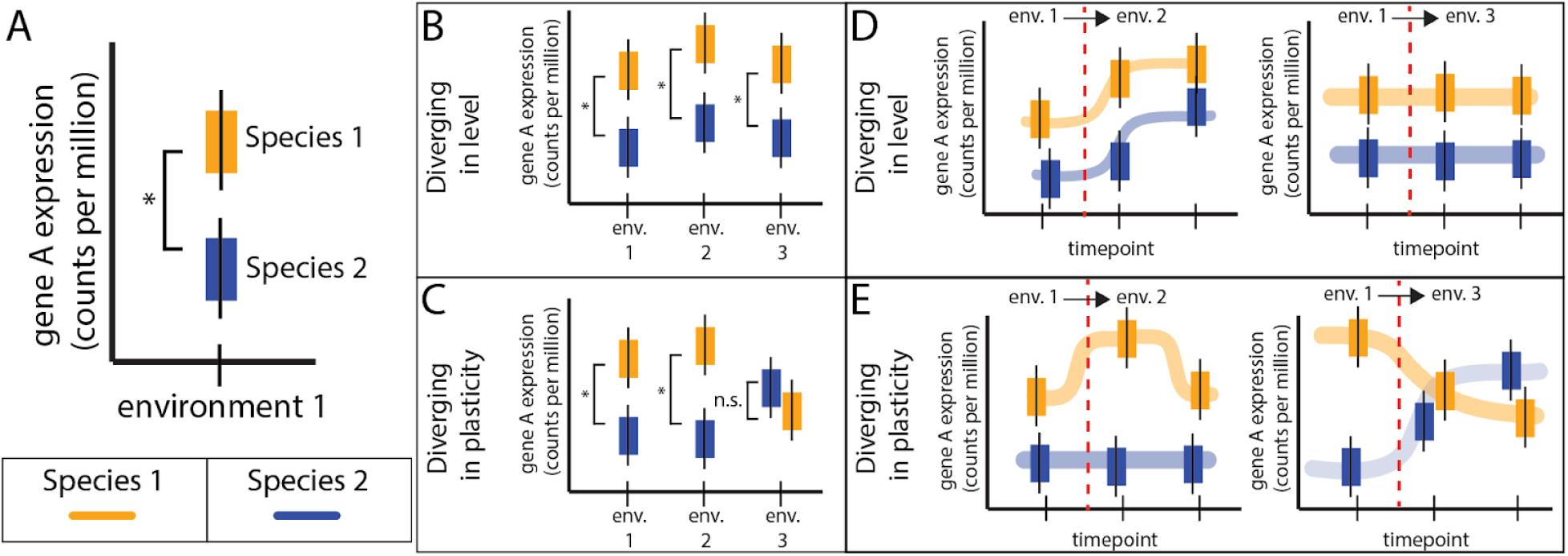
Detecting divergence in gene expression level and plasticity. A) The yellow and blue box plots with error bars show hypothetical results from replicate measures of expression for one gene (Gene A), in two species, in one environment, measured in terms of read counts from Gene A per million read counts. B) The same type of hypothetical data are shown for Gene A, measured in two species, in multiple environments (env. 1, env. 2, env. 3), if the gene’s expression has diverged in a way that is robust to environmental difference (i.e., diverged in expression level). C) Same as in B, but if expression of the gene diverged in an environment-specific manner (i.e., diverged in expression plasticity). D) Two examples are shown of hypothetical expression of Gene A measured in replicate, in two species, at multiple timepoints, transitioning from environment 1 to environment 2 and from environment 2 to environment 3 assuming Gene A diverged only in expression level. E) Same as in D, but if Gene A has diverged in expression plasticity between environments 1 and 2 as well as between environments 1 and 3. In panels D and E, the red dotted line represents the timepoint at which the environment was changed.

Studies contrasting divergence in expression level and expression plasticity have found they are largely evolving independently. For example, in yeast, divergence in mean expression level was found not predict divergence in environment-specific expression patterns (Krieger et al. 2020) A similar independence of divergence in expression level and expression plasticity was reported for separate suites of genes in fish (Dayan et al. 2015). These observations suggest that distinct genetic changes are causing divergence in expression plasticity and overall expression level. Indeed, prior work suggests that *cis*-regulatory divergence tends to be more environmentally robust than *trans-*regulatory divergence (Tirosh et al. 2009; Krieger et al. 2020).

Here, using RNA-seq data from (Krieger et al. 2020) and (Fay et al. 2023), we look more closely at how divergence in gene expression changes when yeast move from one environment to another, comparing expression of orthologous genes in *Saccharomyces cerevisiae* and its close relative *Saccharomyces paradoxus.* These datasets include measures of gene expression from multiple timepoints after transitioning to six different environments. For each environment, we identified divergence in plasticity by using hierarchical clustering to classify each expressed gene in each species as increasing, decreasing, or static following each shift in environment and then looked for differences in the classification of orthologous genes. We found that divergence in plasticity for a given gene was often seen in only one of the six environments. We also found that 12 to 35% of genes examined (depending on the environment) showed evidence of plasticity in one species but not the other, whereas 3 to 27% of genes showed expression increasing in one species but decreasing in the other following the same environmental shift. Consistent with prior work (Krieger et al. 2020), we found that this divergence in plasticity was explained more by *trans*-regulatory divergence than *cis*-regulatory divergence. Finally, using environment-specific regulatory networks inferred using data from Yeastract, we found that divergent plasticity of transcription factors generally did not lead to similar divergence in expression plasticity of their target genes. These differences in divergence between regulators and their target genes could be caused by compensatory changes evolving within the regulatory network and/or robustness to changes in one factor due to the combinatorial control of most genes by multiple transcription factors. Supporting the latter possibility, we found that genes with conserved expression plasticity in a given environment tended to have more regulatory interactions in that environment-specific regulatory network than genes with divergent plasticity. Using these regulatory networks, we also identified five transcription factors in the pleiotropic drug resistance network (YAP1, RPN4, PDR1, PDR3, and YRR1) that had diverged in expression level but showed a higher than typical proportion of target genes with divergent expression plasticity. Taken together, this study advances our understanding of the molecular architecture of divergent gene expression and the regulatory evolution that facilitates ecological differentiation between closely related species.

## Results and Discussion

### Identifying divergence in expression level and expression plasticity

To characterize the extent of divergence in expression level and expression plasticity between *S. cerevisiae* and *S. paradoxus*, we analyzed RNAseq data from both species at multiple timepoints in six different growth conditions (Figure 2): (a) growth to saturation in rich (YPD) media (Krieger et al. 2020), (b) progression through the cell cycle in rich media following synchronization with urea shock (Krieger et al. 2020), (c) transition to and growth in low nitrogen media from rich media (Krieger et al. 2020), (d) transition to and growth in low phosphorous media from rich media (Krieger et al. 2020), (e) transition to and growth at 37C from 25C (heat shock) (Fay et al. 2023), (f) transition to and growth at 12C from 25C (cold shock) (Fay et al. 2023). In each case, total RNA was sampled in the starting condition and then at 3 to 28 timepoints after the change in environment to measure how transcript abundance changed as yeast acclimated to the shift in environment. These data allowed us to determine how the dynamic expression response of yeast cells in multiple environments has diverged between species.

**Figure 2.**
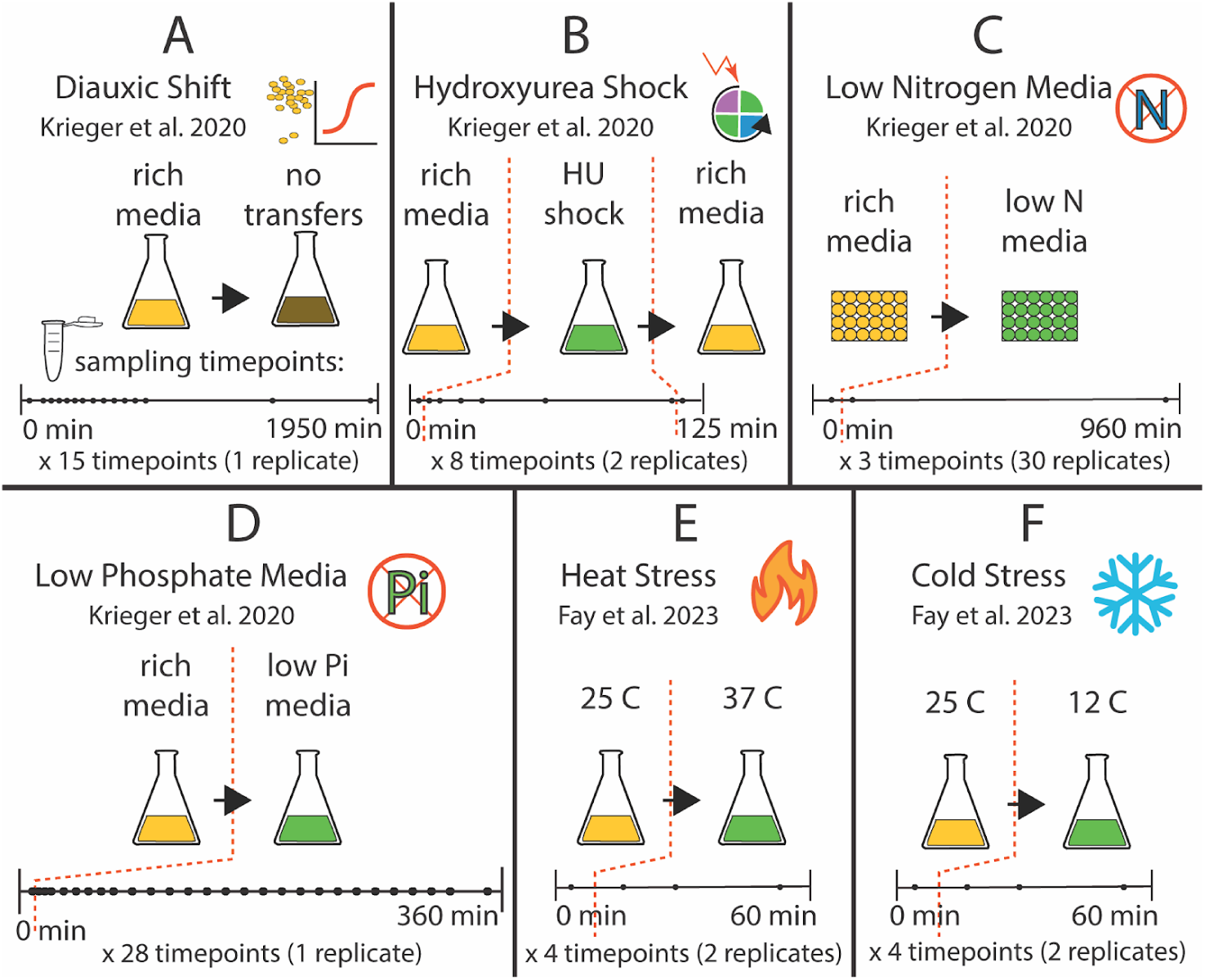
Overview of environmental conditions and sampling. RNA-seq data analyzed was collected from *S. cerevisiae* and *S. paradoxus* in each of the six different growth environments shown, using sampling timepoints and replication shown. A) To capture changes in expression that occurred during the Diauxic Shift, 15 timepoints (black dots on timeline) were sampled with 1 replicate per timepoint. B)-F) Dotted red lines indicate between which two timepoints the yeast were transitioned to a new growth medium. B) To capture changes in expression that occurred in response to hydroxyurea exposure, 8 timepoints were sampled with 2 replicates per timepoint. C) To capture changes in expression that occurred during nitrogen starvation, 3 timepoints were sampled with 30 replicates per timepoint. D) To capture changes in expression that occurred during phosphate starvation, 28 timepoints were sampled with 1 replicate per timepoint. E) To capture changes in expression that occurred during Heat Stress, 4 timepoints were sampled with 2 replicates per timepoint. F) To capture changes in expression that occurred during Cold Stress, 4 timepoints were sampled with 2 replicates per timepoint. The logo shown in the top right of each panel is used to indicate the different environments throughout the paper.

For each of the 165 RNA-seq datasets examined (i.e., each timepoint within each environment with 1-2 replicates per timepoint) (Figure 3A), we first filtered out genes that were lowly expressed in both species (i.e., had average expression less than 30 counts per million), which removed 9-20% of genes in each environment (Figure 3B). Next, we identified genes with significant changes in expression level that were robust to environmental changes (i.e., changes in expression that were seen consistently at all timepoints within an environment). To test for such divergence, we measured the expression level for each gene in each environment by fitting the generalized linear model *Y_ij_ = allele_i_ + timepoint_j_*, where *Y_ij_* is the read count from the RNAseq data for species *i* at timepoint *j* (see Methods). The coefficient for the *allele* term is the estimated log_2_ fold change of *S. cerevisiae*’s read count relative to *S. paradoxus*’, in a single environment, controlling for time-point. This measure of log_2_ fold change describes the average difference in expression level between species independent of any differences in expression plasticity seen as cells transition between environments (Figure 3C). A gene was considered to have diverged in expression level if it had a Wald test adjusted P-value less than 0.05 and a fold change between species greater than 1.5 (log_2_ fold change = 0.585). This adjusted P-value was calculated using the (Benjamini and Hochberg 1995) method to account for multiple testing.

**Figure 3.**
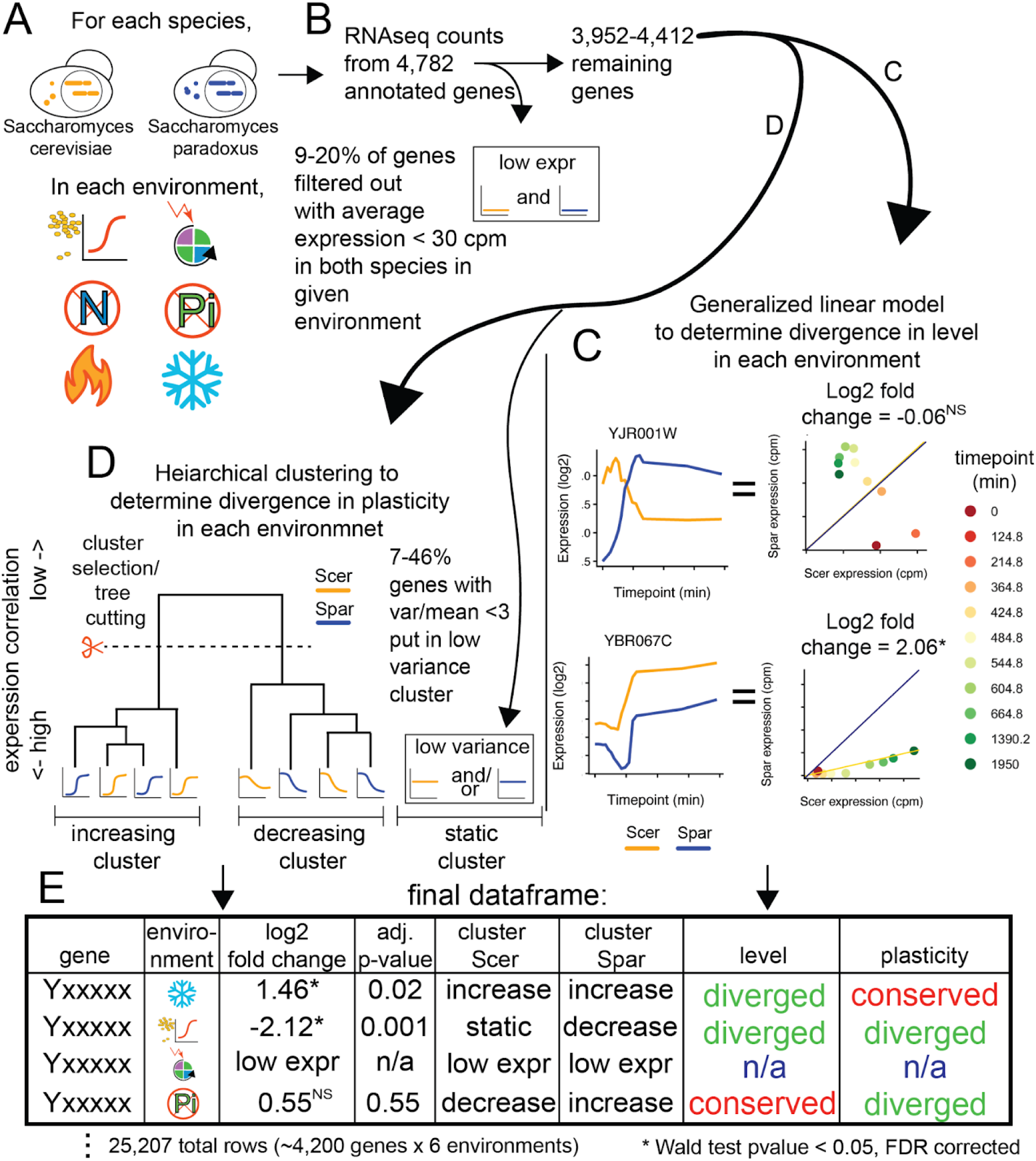
Overview of data analysis. A) The schematic shows a budding cell from *S. cerevisiae* (yellow) and *S. paradoxus* (blue) along with the six different environments analyzed, represented by the logos defined in Figure 2. B) A subset of genes were excluded for being lowly expressed in both species in the focal environment (9-20% of genes, depending on the environment). C) The left two panels show the expression (log_2_ fold change) of two genes (YJR001W and YBR067C) during growth to saturation in YPD (capturing the diauxic shift) for *S. cerevisiae* (Scer, yellow) and *S. paradoxus* (Spar, blue). The right two panels compare the expression (measured in counts per million reads, or cpm) of each of these genes between *S. cerevisiae* (Scer expression) and *S. paradoxus* (Spar expression), with the data from different time points shown in different colors. A black Y=X line is shown for reference on each plot. YJR001W (top) provides an example of a gene without a consistent difference in expression between species across all timepoints in this environment (indicated by “NS”, reflecting a Wald test adjusted p-value > 0.05). YBR067C (bottom) shows an example of a gene with a consistent expression difference across all timepoints in this environment, with the general linear model estimating the average log_2_ fold change difference across all timepoints to be 2.06, with a Wald test adjusted p-value < 0.05, as indicated by the asterisk. D) Measuring divergence in expression plasticity for five pairs of orthologous genes (ten genes total). Two genes had a variance/mean ratio (dispersion) less than 3, so were placed in the low variance (“static”) cluster. A hypothetical gene tree resulting from hierarchical clustering of the remaining 8 genes is shown, with four genes assigned to the “increasing” cluster and four assigned to the “decreasing” cluster. E) Data and inferences from these analyses are summarized in Table S1. The figure shows hypothetical data in the structure of Table S1 as described in the main text to illustrate the different types of expression divergence. Note that the “adj. P-value” column shows the result of a Wald test from the linear model testing for divergence in expression level with a Benjamini Hochberg correction for multiple testing, which determines whether the “level” is inferred to be “conserved” or “diverged”. “Plasticity” is inferred to be conserved if “cluster Scer” and “cluster Spar” have the same value (except when the gene is lowly expressed in both species).

To measure expression plasticity (i.e., differences in expression level specific to a subset of timepoints within an environment), we grouped the expression levels of all genes in both species at all timepoints by hierarchical correlation clustering (Figure 3D). This approach is similar to that used in the R package WGCNA (Langfelder and Horvath 2008). This method produces a hierarchical dendrogram (or “tree”) based on expression correlations that includes all genes in both species. Genes downstream of a branch “cut” in this tree are considered to be in the same cluster (Figure 3D). Genes in the same cluster have more highly correlated expression profiles than those in different clusters. The number of clusters identified from this analysis is determined by where the user chooses to cut this tree (Figure 3D). Prior to clustering, we assigned genes that had low expression variability among timepoints to a “low-variance cluster,” as these genes were weakly correlated with all other genes (Figure 3D). Depending on the environment analyzed, this low-variance cluster included 7 to 46% of genes. After clustering, we identified genes diverging in plasticity between species by identifying cases where one-to-one orthologs were placed in different clusters (Figure 3D). We note that correlation clustering is independent of any constitutive changes in expression level because correlations are invariant to multiplying by a constant (e.g., log_2_ fold change).

To determine how the choice of cut point would affect our analysis, we compared the clusters of genes resulting from cut points that generated 2, 3, or 4 clusters. We found that 97% of the genes identified as having orthologs in different clusters in the 2-cluster scheme were also identified as having orthologs in different clusters in the 3- and 4-cluster schemes (Figure S1). That is, nearly all genes that we interpret as having divergent plasticity with the 2-cluster scheme are also interpreted as having divergent plasticity with the 3- or 4-cluster schemes. If we consider the converse, about 80% of the genes identified as having orthologs in different clusters in the 3- or 4-cluster scheme were also identified as having orthologs in different clusters in the 2-cluster scheme, indicating that the 2 cluster scheme is more conservative for identifying genes with divergent plasticity. We thus chose to focus our analysis on the 2-cluster scheme. Looking at the expression of genes in the two clusters within each environment showed that genes in one cluster tended to increase their expression following the shift in environmental conditions, whereas genes in the other cluster tended to decrease their expression at that time. We thus classified each gene in each species as having “increased” expression, “decreased” expression, or “static” expression (if the gene met the criteria for inclusion in the “low variance” set).

To better understand the expression profiles of genes classified as having increased, decreased, or static expression over time within each environment, we plotted the mean expression level (log_2_ counts per million) of all genes in each category at each timepoint in each environment (Figure 4). We noted that the increasing and decreasing clusters showed different patterns of average expression changes among environments, with heat and cold stress being the only environments not to show monotonic changes in average expression over time (Figure 4). We also noted that the time course of average expression levels of genes in the increasing and decreasing clusters were largely symmetrical; we understand this to be because the hierarchical correlation clustering maximizes the positive correlation among genes within a group, which also maximizes the negative correlation between the two groups. In all six environments, the majority of genes (79%, 88%, 72%, 93%, 76%, 54%) showed variable expression (increased or decreased during the transition) rather than static expression. The highest proportion of plastic genes was observed in the low phosphate environment; only 625 of 8766 (7%) genes were classified as having static expression (Figure 4). The lowest proportion of plastic genes was observed in the cold stress environment; 3733 of 8102 genes (46%) were classified as having static expression (Figure 4). The average expression level of genes in the static cluster (i.e., genes expressed consistently among timepoints) was consistently lower than the average expression level of genes with variable expression among timepoints (Figure 4), suggesting that genes exhibiting expression plasticity tend to be expressed at higher average expression levels than genes that do not.

**Figure 4.**
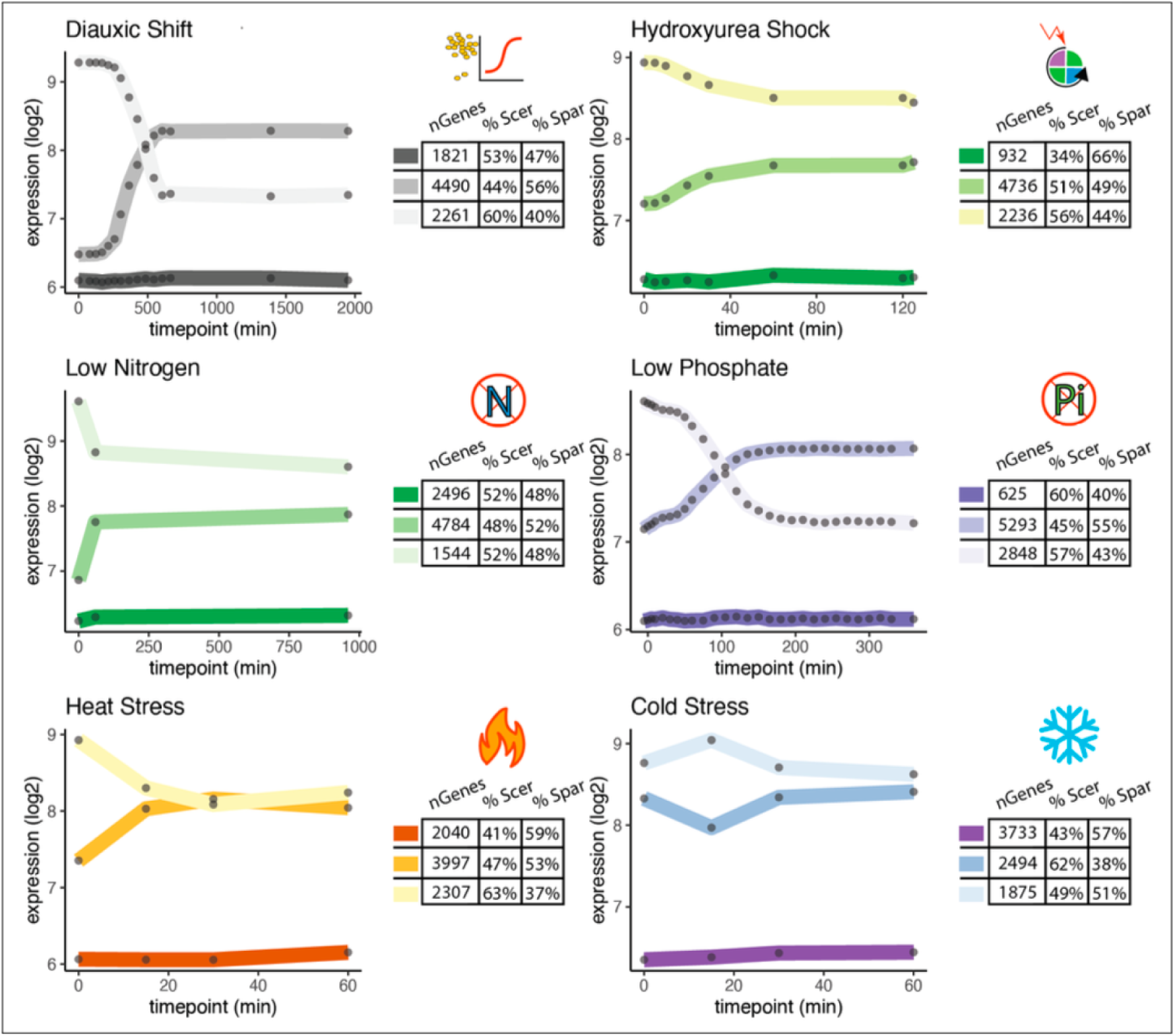
Contrasting average expression of genes classified as increasing, decreasing, or static. In each panel, the lines show the average expression of all genes in each cluster, with black dots on the lines showing the timepoints at which samples were taken. The number of genes in each cluster (nGenes, counting *S. cerevisiae* and *S. paradoxus* orthologs separately) is shown in the table to the right of each line plot. The percentages of these genes from *S. cerevisiae* (% Scer) versus *S. paradoxus* (% Spar) are shown beside the overall gene counts. Logos represent the six different environments, as defined in Figure 2.

We generated a data table (Figure 3E, Table S1) containing one row for each of the ∼4,200 genes with a one-to-one ortholog in *S. cerevisiae* and *S. paradoxus* in each of the 6 environments This table includes the gene name; the environment; the log_2_ fold change reported by the general linear model representing the average expression difference between species across all timepoints within the environment; the statistical significance of that fold change based on a Wald test, adjusted for multiple testing; whether the *S. cerevisiae* ortholog was in the increased, decreased, or static cluster or was excluded from analysis because it was lowly expressed in both species; and the same information for the *S. paradoxus* ortholog. Based on this information, we recorded whether each gene showed evidence of divergence in expression level and/or expression plasticity. As described above, genes with a statistically significant Wald test and a log_2_ fold change expression difference greater than 0.585 were considered to be diverged in terms of expression level, and genes with *S. cerevisiae* and *S. paradoxus* orthologs assigned to different correlation clusters (i.e., increase, decrease, or static) were considered to have divergent expression plasticity (Figure 3E, Table S1).

### Divergence in expression as S. cerevisiae and S. paradoxus go through the diauxic shift

To better understand divergence in expression level and expression plasticity between *S. cerevisiae* and *S. paradoxus* at the level of individual genes, we focused first on expression changes during the diauxic shift. (Parallel analyses were performed for the other five environments, and the results are shown in Figure S2.) When *S. cerevisiae* or *S. paradoxus* are grown in media with glucose as a carbon source (such as Yeast Extract Peptone Dextrose, or YPD), the cells initially ferment glucose. When the glucose is depleted, they undergo a significant metabolic change (the diauxic shift) and begin respiring using the ethanol produced by fermentation. This transition is well-characterized in *S. cerevisiae* and known to involve many changes in gene expression (Galdieri et al. 2010; Hossain et al. 2016; Cha et al. 2021).

We found that 577 genes were lowly expressed (< 30 counts per million) at all timepoints in both species during growth to saturation in YPD media, leaving 4286 one-to-one ortholog pairs for analysis in each species, or 2 x 4286 = 8572 total genes. In the diauxic shift environment, the static cluster consisted of 1821 total genes, with 971 belonging to *S. cerevisiae* and 850 belonging to *S. paradoxus*. The increasing cluster consisted of 4490 genes, with 1962 belonging to *S. cerevisiae* and 2516 belonging to *S. paradoxus*. The decreasing cluster consisted of 2261 genes, with 1353 belonging to *S. cerevisiae* and 920 belonging to *S. paradoxus* (Figure 4).

These differences between species in the number of genes belonging to each cluster suggest that the regulation of genes has diverged during the diauxic shift. Based on the tests for conservation and divergence of expression level and expression plasticity described above (Figure 3), we found that 49% of genes (2089/4286) had both conserved expression level and conserved expression plasticity between *S. cerevisiae* and *S. paradoxus* (Figure 5A). This group includes genes where the *S. cerevisiae* and *S. paradoxus* orthologs were both in the increasing cluster, both in the decreasing cluster, or both in the static cluster. The mean expression level is shown in Figure 5B for the 1218 genes for which both orthologs were in the increasing expression cluster. 17% of genes (742/4286) had diverged in mean expression level but not expression plasticity (Figure 5A). In such cases, we detected a significant difference in mean expression between species with the Wald test, but both orthologs were in the same plasticity cluster. Figure 5C shows an example of this pattern for mean expression of the 284 genes in the increasing cluster. 24% of genes (1015/4286) had a conserved mean expression level but divergent expression plasticity (Figure 5A). To provide an example of this category, the mean expression is shown for the 188 genes with the *S. cerevisiae* ortholog in the decreasing category and the *S. paradoxus* ortholog in the increasing category in Figure 5D. Finally, 10% of genes (440/4286) showed evidence of divergence in both expression level and expression plasticity (Figure 5A), which is illustrated in Figure 5E by showing mean expression for the 122 genes categorized as having static expression in *S. cerevisiae* and increasing expression in *S. paradoxus*, with the higher expression level in *S. paradoxus*.

**Figure 5.**
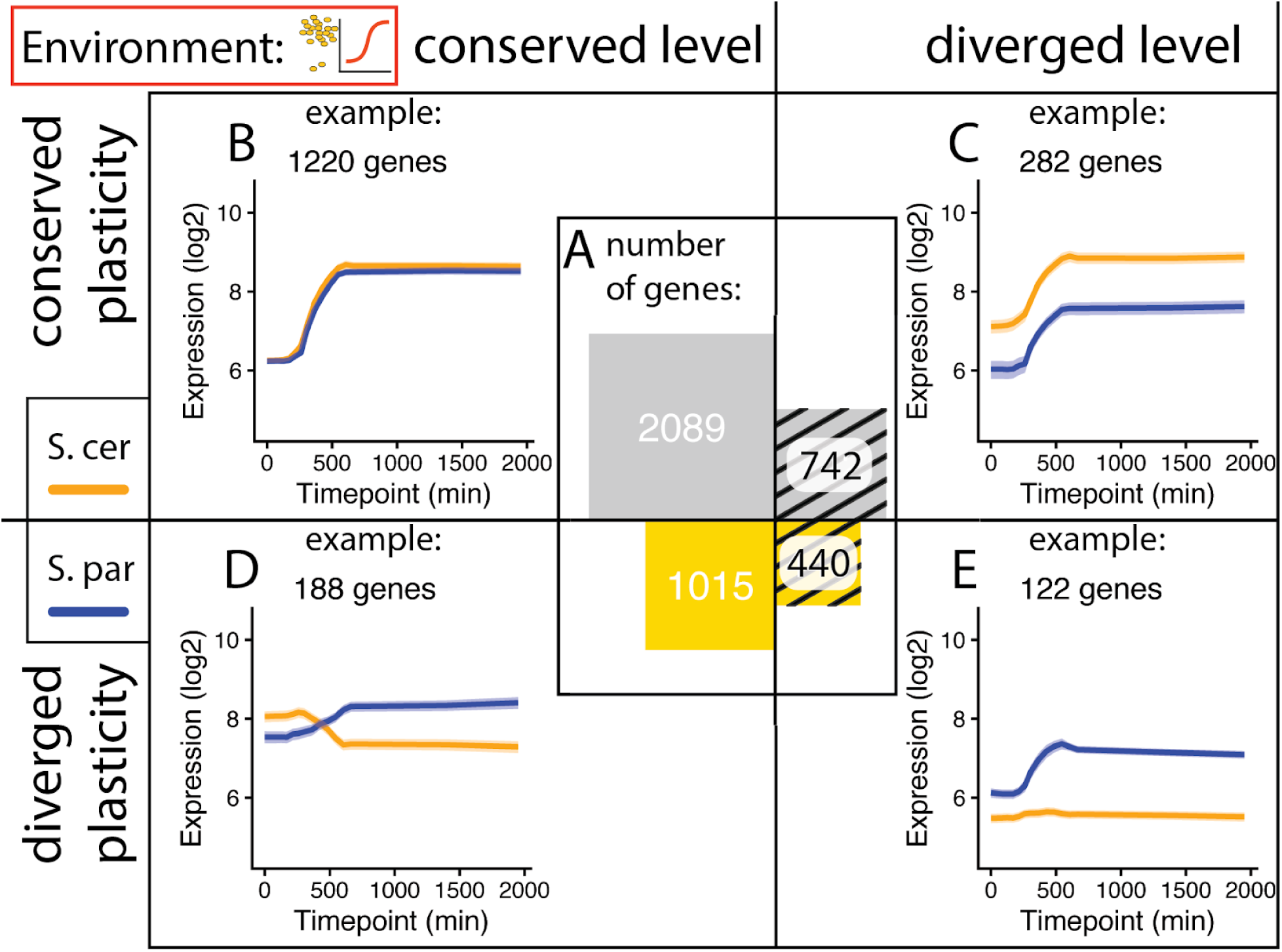
Separating divergence in expression level from divergence in expression plasticity in the Diauxic Shift environment. A) Proportional area plot shows the total number of genes in each of the 4 divergence categories. B) Average expression of orthologs with conserved expression level and plasticity that increase expression at the diauxic shift is shown for *S. cerevisiae* (S. cer) in yellow and *S. paradoxus* (S. par) in blue. C) Average expression of orthologs that have a higher expression level in *S. cerevisiae* and increase expression at the diauxic shift. D) Average expression of orthologs that increase expression in *S. paradoxus* and decrease expression in *S. cerevisiae* at the diauxic shift. E) Average expression of orthologs that have a higher expression level in *S. paradoxus* and increase expression at the diauxic shift only in *S. paradoxus*. In panels B-D, shading around the lines are error ribbons that indicate the 95% confidence interval for the mean.

The relative number of genes in each category is consistent with prior findings that changes in expression level and plasticity tend to evolve independently (Krieger et al. 2020), although there were more genes diverging in both expression level and plasticity in this dataset focused on the diauxic shift than expected by chance (Fisher Exact Test odds ratio 1.22, p-value < 0.001). Similar tests for independence using the data from the other five environments (Figure S2) showed fewer genes diverging in both level and plasticity than expected in the heat stress environment (Fisher exact test odds ratio 0.95, p-value < 0.04), with no significant relationship observed in any of the other 4 experiments.

### Similar portions of genes have divergent expression plasticity among environments

To determine whether the expression divergence seen in the diauxic shift environment was typical, we compared it to the expression divergence seen in the other five environments. We found that the proportions of genes that were conserved or divergent in gene expression level and/or plasticity were similar in all six environments (Figure 6A). In all cases, the largest portion of genes showed conserved level and plasticity (solid grey in Figure 6A), roughly equal numbers of genes had diverged only in level (striped grey in Figure 6A) or only in plasticity (solid gold in Figure 6A), and the smallest portion of genes had diverged in both level and plasticity (striped gold in Figure 6A). Increasing the fold change cutoff used to classify genes as having divergent expression from 1.5 to 2 changed the frequency of genes in these categories but retained similar patterns among the six environments, whereas eliminating the fold change cutoff (i.e., relying only on significance of the Wald test) caused more significant differences in these patterns among environments (Figure S3).

**Figure 6.**
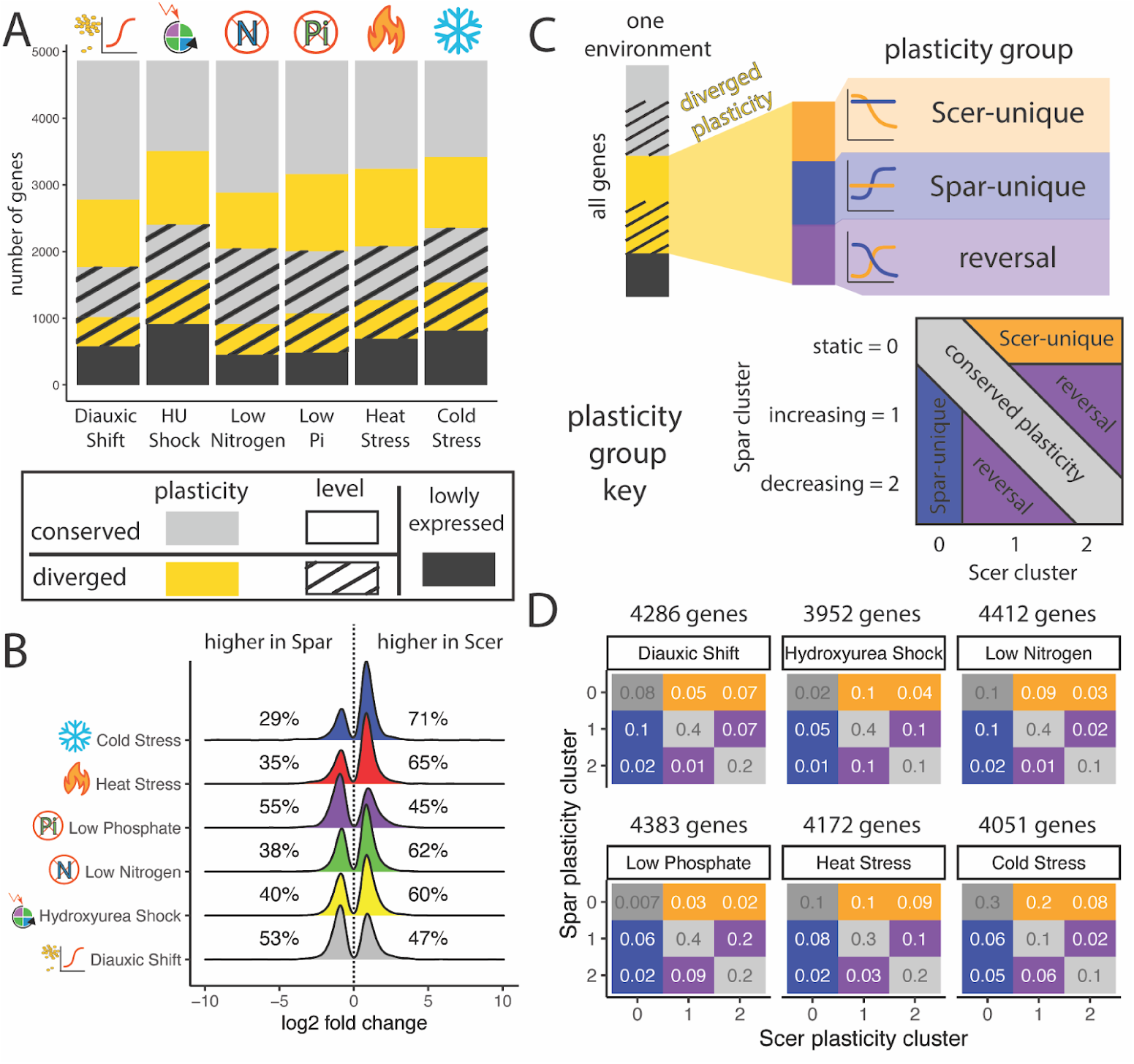
Similar portions of genes are diverging in level and plasticity in each environment. A) Stacked barplot shows the number of genes diverging in expression level (hatched) and expression plasticity (yellow) between *S. cerevisiae* and *S. paradoxus* in each environment. Logos representing environmental changes are defined in Figure 2. B) Ridgeline plot showing the distribution of log_2_ fold changes in gene expression for genes with significant divergence in expression level in each environment. C) Categories of divergence in gene expression plasticity. “Scer-unique” and “Spar-unique” include genes classified as plastic in *S. cerevisiae* or *S. paradoxus,* respectively, and static in the other species. “Reversal” includes genes that are increasing expression after shifting to a new environment in one species but decreasing after the same expression shift in the other species. The plasticity group key shows how the cluster (static, increasing, or decreasing) containing the *S. paradoxus* (Spar) and *S. cerevisiae* (Scer) orthologs relates to these three categories as well as a fourth category of conserved expression plasticity. D) For each of the six environments, the proportion of genes with orthologs in each combination of plasticity clusters is shown. The number of orthologous gene pairs analyzed in each environment is shown above each matrix. Boxes are color-coded based on the plasticity group key shown in panel C.

Considering only genes with significant divergence in expression level in each environment, we found that the *S. cerevisiae* ortholog was more highly expressed for the majority of genes in the cold stress, heat stress, low nitrogen, and hydroxyurea shock environments (Figure 6B). The *S. paradoxus* ortholog was more highly expressed in the majority of genes in the low phosphate and diauxic shift environments (Figure 6B). In all environments, at least 45% of genes had higher expression in *S. cerevisiae*. In other words, we observed no consistent shift in expression divergence toward higher expression in one species or the other among environments.

For genes with significant divergence in expression plasticity, we considered that a gene’s expression could be plastic (i.e., show significant changes in expression among timepoints) in one species but static in the other (i.e., plasticity is unique to *S. cerevisiae* or *S. paradoxus*), or that both species could show significant plasticity but with changes in expression in opposite directions (i.e., a reversal, with expression increasing in one species and decreasing in the other) (Figure 6C). To identify genes in each of these groups, we examined each pair of orthologous genes and noted whether the *S. cerevisiae* and *S. paradoxus* orthologs were in the static, increasing, or decreasing cluster. Genes with the *S. cerevisiae* ortholog in the static cluster but the *S. paradoxus* ortholog in the increasing or decreasing cluster were considered to have plasticity unique to *S. paradoxus*, and vice versa. Genes with *S. cerevisiae* ortholog in the increasing cluster and the *S. paradoxus* ortholog in the decreasing cluster (and vice versa) were considered to show a reversal in plasticity (Figure 6C). Genes with both orthologs in the same plasticity cluster (static, increasing, or decreasing) were considered to have conserved plasticity (Figure 6C).

As expected, conserved plasticity was the most common of these categories in all environments: 55 to 70% of the 3952 to 4412 genes analyzed in each environment had both orthologs in the same plasticity cluster (Figure 6D). Species-specific plasticity, in which one ortholog was considered plastic (increasing or decreasing) and the other was considered static, was observed for 12 to 35% of genes, depending on the environment. Finally, 3 to 27% of genes showed evidence of reversals between species in the direction of expression change after moving to a new environment, depending on the environment (Figure 6D). Reversals in plasticity between species were particularly common following hydroxyurea treatment (25% of genes) and transfer to a low phosphate environment (27% of genes), suggesting that *S. cerevisiae* and *S. paradoxus* might have diverged the most in terms of how they respond to these particular environmental shifts.

We found reversals in the direction of expression plasticity to be particularly surprising and wondered how the threshold used to classify genes in the static expression category affected their frequency. We therefore repeated this analysis with a higher (var/mean = 5) and lower (var/mean = 1) threshold for considering a gene to have static expression than that used for our primary analysis (var/mean = 3) (Figure S4). Even with the threshold that is most conservative for identifying genes as plastic (i.e., var/mean = 5), 2% to 16% of genes still showed a reversal in the direction of expression change as cells acclimate to the new environment (Figure S4). The molecular mechanisms responsible for such changes in plasticity remain unknown, but a transcription factor changing from an activator to a repressor in a context-dependent manner could explain these observations. Many transcription factors that can both activate and repress gene expression have been identified in *S. cerevisiae*, including Rap1p (Shore and Nasmyth 1987) and Ume6p (Raithatha et al. 2021).

### Divergence in expression plasticity tends to be environment-specific

With the transition to all six environments showing similar proportions of genes diverged in expression level and expression plasticity between *S. cerevisiae* and *S. paradoxus*, we asked whether the same genes tended to show similar changes in expression level and expression plasticity among the six environments. We first assessed this question by looking at whether the same gene tended to appear in the same expression divergence category in multiple environments. As shown in Figure 7A, we found that the divergence category in the diauxic shift environment was qualitatively a poor predictor of the divergence category in other environments. We then compared expression divergence among environments more quantitatively by (a) identifying genes with a particular pattern of expression divergence in one environment and then (b) examining expression of that set of genes in all six environments. We performed this analysis for both expression level (measured as the average log_2_ fold change) and expression plasticity (measured as the average correlation in expression between the *S. cerevisiae* and *S. paradoxus* orthologs following an environmental shift), as shown in Figures 7B and C, respectively. For expression level, genes with higher expression in *S. cerevisiae* were analyzed separately from genes with higher expression in *S. paradoxus* (Figures 7D and E). Similarly, genes with expression increasing in *S. cerevisiae* and decreasing in *S. paradoxus* after shifting to a new environment were analyzed separately from genes increasing in *S. paradoxus* and decreasing in *S. cerevisiae* (Figure 7F). Genes with static expression in either species were excluded from this analysis because low correlations are expected with any other expression pattern. Consequently, only genes classified as having reversals in plasticity between species (Figure 6D) were included in the analysis shown in Figure 7F.

**Figure 7.**
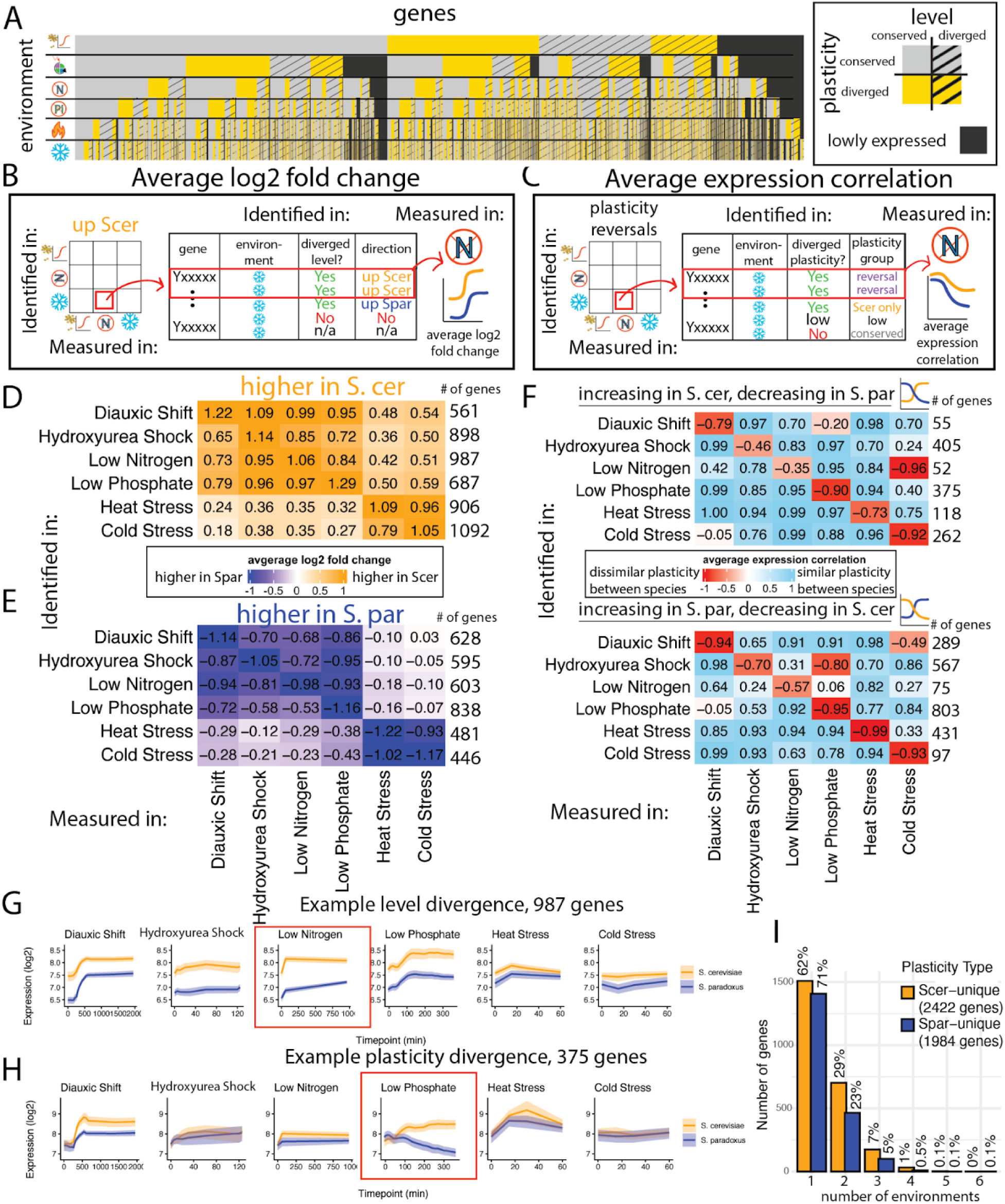
Divergence in expression plasticity tends to be environment-specific. A) Categorical heatmap of divergence category for each gene in each environment. Genes are ordered by their category in the environment listed in the top row first, followed by their category in the second, third, fourth, fifth, and six rows in a nested fashion. B-C) Schematics show how genes were selected for analysis based on their divergence in expression level (B) or expression plasticity (C) in one environment (cold stress, in this case) and then analyzed (“measured in”) each of the six environments (with low nitrogen shown as an example environment). They also show how the results of these analyses (average log2 fold change for gene expression level in B and average expression correlation for expression plasticity in C) are reported in the matrices of panels D, E, and F. In (B), “up Scer” and “up Spar” refer to genes with higher expression in *S. cerevisiae* or *S. paradoxus,* respectively. In (C), “Scer only” refers to genes with plastic expression in *S. cerevisiae* and static expression in *S. paradoxus*. Only genes classified as reversals (i.e., expression increasing in *S. cerevisiae* and decreasing in *S. paradoxus* or vice versa) were included in the analysis shown in panel F. D-E) Heatmaps show the average log_2_ fold change in expression for sets of orthologous genes identified as having a divergent expression level in the environment shown to the left of the matrix, measured in the environment shown at the bottom of the matrix, for genes with higher expression in *S. cerevisiae* (D) or *S. paradoxus* (E). The number to the right of each row is the number of genes included in each analysis shown in that row. F) Heatmaps show the expression correlation for orthologous genes with a reversal in the direction of plasticity in the environment shown on the left of the matrix, in the environment shown at the bottom of the matrix. Genes with expression increasing in *S. cerevisiae* are shown in the top matrix and genes with expression decreasing in *S. cerevisiae* are shown in the bottom matrix. G-H) The average expression of the set of genes with divergent expression level (G) or divergent expression plasticity with a reversal of direction (H) is shown in each of the six environments for the set of genes selected based on data from the low nitrogen environment for expression level (G) or low phosphate environment for expression plasticity (H). Red boxes indicate the environment the groups of genes were identified in. The yellow line shows average expression of *S. cerevisiae* orthologs and the blue line shows average expression of the *S. paradoxus* orthologs for genes in each of these gene sets, with shading around the line showing the 95% confidence intervals for the mean. I) For the 2422 genes with plastic expression in *S. cerevisiae* and static expression in *S. paradoxus* (yellow), as well as the 1984 genes that showed the opposite pattern (blue), the number of genes showing that pattern of expression in 1, 2, 3, 4, 5, or all 6 environments is shown.

We found that genes with a divergent expression level and higher expression in *S. cerevisiae* (positive log_2_ fold change values) in one environment also tended to have a higher expression level in *S. cerevisiae* in all five other environments (Figure 7D). In each case, the largest average expression difference between *S. cerevisiae* and *S. paradoxus* was seen in the environment used to identify the set of genes as having divergent expression levels (i.e., values along the diagonal in Figure 7D). This pattern is further illustrated in Figure 7G, which shows the average expression level in each species, in each of the six environments, for the set of 987 genes identified as having a divergent expression level with higher expression in *S. cerevisiae* in the low nitrogen environment.

Genes identified in each environment as having a divergent expression level with higher expression in *S. paradoxus* (negative log_2_ fold change values) showed the same patterns. The average difference in expression level for a set of genes was greatest between species in the environment used to define that set of genes (i.e., values along the diagonal in Figure 7E), and genes identified as having divergent expression with higher expression of the *S. paradoxus* ortholog in one environment tended to also have higher expression of the *S. paradoxus* ortholog in other environments. The one exception was 628 genes identified as divergent in the diauxic shift environment, which showed an average log_2_ fold change in expression level of 0.02 (i.e., very similar average expression level between species) in the cold stress environment (Figure 7E).

The analyses shown in Figures 7D and 7E also revealed that the magnitude of average differences in expression level between species tended to be more similar among the diauxic shift, hydroxyurea shock, low nitrogen, and low phosphate environments than the heat and cold stress environments. While these differences might result from how yeast respond to these different types of stresses, they could also result from the fact that data was collected for these two groups of environments by different labs, using different strategies for RNA-seq, and included different strains of *S. cerevisiae* (S288c vs. T73) and *S. paradoxus* (CBS432 vs. N17) (see Methods). We note, however, that despite these differences in magnitude, these sets of environments do not differ in the direction of the expression differences between species. Observing consistent directionality in the average changes in expression levels among environments suggests that the genetic changes that cause one ortholog to be more highly expressed than the other ortholog among timepoints in a single environment also tend to have consistent effects on expression level among environments.

For expression plasticity, we found that orthologs classified as having a reversal of plasticity in one environment (negative correlation coefficient) tended to only show a negative correlation in that environment (values along the diagonals in Figure 7F). In most other environments, these sets of genes tended to have a positive correlation (indicating conserved plasticity) between expression of the *S. cerevisiae* and *S. paradoxus* orthologs. An example of this pattern is shown in Figure 7H, where the average expression of the 375 genes showing evidence of a reversal in plasticity between the *S. cerevisiae* and *S. paradoxus* orthologs in low phosphate is shown for all six environments. Note the conserved expression between orthologs in the time course for all environments except low phosphate (Figure 7H). This observation suggests that the genetic differences between *S. cerevisiae* and *S. paradoxus* responsible for divergence in expression plasticity tend to have environment-specific effects.

Notable exceptions to this pattern of environment-specific plasticity divergence include the set of 52 genes that showed a reversal of plasticity in low nitrogen with expression increasing in *S. cerevisiae* (top matrix in Figure 7D), but also showed an even stronger negative correlation in expression in the cold stress environment (r = -0.35 in low nitrogen, r = -0.96 in cold stress). Similarly, the 567 genes identified as having a reversal in the direction of plasticity between species following hydroxyurea shock with expression increasing in *S. paradoxus* (bottom matrix in Figure 7D) showed an even more negative correlation following transfer to low phosphate media (r = -0.70 in hydroxyurea shock, r = -0.80 in low phosphate). Finally, the 289 genes showing a reversal in plasticity in the diauxic shift experiment for genes decreasing in expression over experimental time in *S. paradoxus* also showed a negative correlation between orthologs in the cold stress environment (r = - 0.94 in diauxic shift, r = -0.49 in cold stress). For these three comparisons, Figure S5 shows the average expression of these sets of genes in each species, in all six environments, similar to Figure 7H. It will be interesting to explore in future work whether these exceptions reflect divergence in transcription factors (or other regulators) that are shared between these pairs of environments for these gene sets.

Finally, we assessed the environmental specificity of genes classified as showing expression plasticity (increasing or decreasing) in one species but static expression in the other species (Scer-unique and Spar-unique in Figures 6C and D). We did this by looking at the 2422 genes classified as Scer-unique in at least one environment and asking how many environments of the six total environments showed a Scer-unique pattern. We also looked at the 1984 genes classified as having Spar-unique plasticity in at least one environment and asked the same question. We found that 62% and 71% of genes, respectively, showed plasticity only in *S. cerevisiae* or *S. paradoxus*, respectively, that was limited to only one of the six environments (Figure 7I), consistent with the divergence of expression plasticity tending to be environment-specific.

### trans-regulatory changes tend to be responsible for divergence in expression plasticity

To examine the molecular mechanisms responsible for the divergence in expression level and expression plasticity, we used RNA-seq data from F_1_ hybrids between *S. cerevisiae* and *S. paradoxus* that was collected in parallel with the RNA-seq data from each of the parental species in each environment (Krieger et al. 2020; Fay et al. 2023). Differences in expression seen between the two species that are also seen between the two alleles in F_1_ hybrids are attributable to *cis*-regulatory divergence (e.g., changes in promoter sequences), whereas differences observed between the two species that are not recapitulated in allele-specific expression of F_1_ hybrids are attributable to *trans*-regulatory divergence (e.g., changes in transcription factors or other diffusible molecules that impact transcription) (Wittkopp et al. 2004).

To estimate the proportion of divergence in expression level attributable to *cis-* or *trans*-regulatory divergence, we compared the average log_2_ fold changes among timepoints between parental species with the magnitude of average log_2_ fold changes between hybrid alleles. To estimate the proportion of divergence in expression plasticity attributable to *cis-* or *trans*-regulatory divergence, we compared the expression correlation among timepoints between *S. cerevisiae* and *S. paradoxus* to the expression correlation between hybrid alleles. We also examined the similarity in plasticity by looking at the scaled mean difference in expression among timepoints. The log_2_ fold changes, expression correlations, and scaled mean differences used in this section are summarized in Table S2.

Depending on the type of expression divergence (level or plasticity) and the mode of expression divergence (*cis* or *trans*), a range of scenarios are possible for each gene in each environment. For example, if we observe a gene with a large log_2_ fold change between parental species and a similarly large log_2_ fold change between hybrid alleles, we would conclude that the genetic change(s) responsible for this divergence in expression level act primarily in *cis* (Figure 8A). If instead we observe a large log_2_ fold change in magnitude between parents and a small log_2_ fold change between hybrid alleles, we would conclude that the genetic change(s) responsible for this divergence in expression level act primarily in *trans* (Figure 8A). Similarly, if a gene shows a weak or negative correlation between parental alleles and a similarly weak or negative correlation between hybrid alleles, we would conclude that the genetic change(s) responsible for this divergence in plasticity act primarily in *cis* (Figure 8A). If we instead observe genes with a weak or negative correlation between parental alleles and a strongly positive correlation between hybrid alleles, we would conclude that the genetic change(s) responsible for this divergence in expression plasticity act primarily in *trans* (Figure 8A). These types of inferences become more complex when both *cis-* and *trans*-regulatory changes contribute to divergent expression level and/or plasticity of the same gene. However, one clear case is when genes have little to no expression divergence between species but show differences in expression between the species-specific alleles in F_1_ hybrids, which indicates the presence of both *cis*- and *trans*-regulatory changes with compensatory effects (Figure 8A).

**Figure 8.**
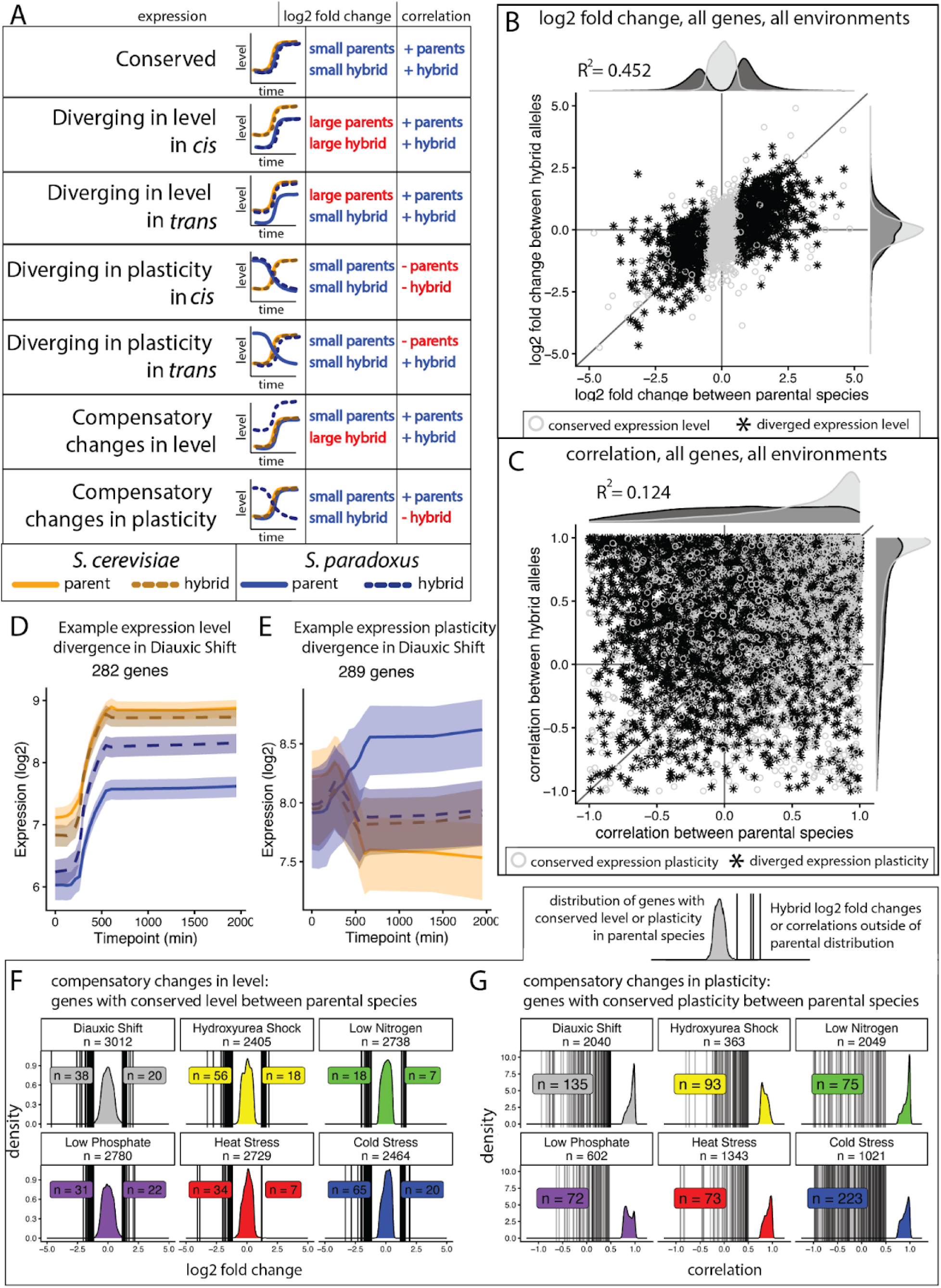
Contributions of *cis*- and *trans*-regulatory divergence to divergence in expression level and plasticity. A) Schematic shows patterns of gene expression divergence expected between species and between alleles in F_1_ hybrids depending on whether the genetic variation responsible for expression divergence acts in *cis* or in *trans*. Plots show hypothetical data that would be consistent with the type of regulatory divergence shown in each row. The relative differences in expression level (small or large) and sign (+ or -) of a correlation in expression between the two species (parents) or between the two alleles in F_1_ hybrids (hybrid) expected in each case are also shown. B) The scatterplot compares the log_2_ fold change in expression between the parental species (*S. cerevisiae* and *S. paradoxus*) to the log_2_ fold change in expression between alleles in the F_1_ hybrids. Each point is one gene in one environment. Black asterisks show data for genes identified as having divergent expression level between parental species, whereas the open grey circles show data for genes classified as having conserved expression between species (based on the Wald test P-value and a log_2_ fold change threshold of 0.585). C) The scatterplot compares the expression correlation between parental species (*S. cerevisiae* and *S. paradoxus*) to the expression correlation between species-specific alleles in F_1_ hybrids. Each point is one gene in one environment. Black asterisks show data for genes identified as having divergent expression plasticity between species, whereas the open grey circles show data for genes identified as having conserved expression plasticity. D) Average expression is shown for the 284 genes identified as having diverged in expression level with higher expression level in *S. cerevisiae* in the Diauxic Shift environment. The solid yellow and blue lines represent expression in *S. cerevisiae* and *S. paradoxus*, respectively, and the dotted yellow and blue lines represent expression of the S.cerevisae and S paradoxus alleles in F_1_ hybrids. The shaded areas indicate the 95% confidence intervals of the mean. E) Average expression is shown for the 289 genes identified as having divergent expression plasticity with expression decreasing in *S. cerevisiae* and increasing in *S. paradoxus* in the Diauxic Shift environment. F) For each environment, the distribution of log_2_ fold change in expression between species is shown for genes classified as having conserved expression level (Wald p-value > 0.05 and fold change in expression < 1.5, which is a log_2_ fold change < 0.585). Despite this similar expression between species, some genes showed large differences in expression between species-specific alleles in F_1_ hybrids that suggest compensatory *cis*- and *trans*-regulatory divergence. The vertical lines shown on each plot indicate the log_2_ fold change in expression between alleles in the F_1_ hybrids for genes where the fold change was greater than 3 (log_2_ fold change > 1.585). The numbers of such genes with higher expression of the *S. cerevisiae* allele or the *S. paradoxus* allele in F1 hybrids are shown on the plot for each environment. G) The distribution of correlation coefficients between *S. cerevisiae* and *S. paradoxus* is shown for genes considered to have conserved expression plasticity and an expression correlation > 0.75 for each environment. Vertical lines show the expression correlation between species-specific alleles in the F_1_ hybrids for genes where this correlation was < 0.5. The number of such genes is shown for each environment. These genes represent cases where *cis*-regulatory divergence causes differences in expression plasticity between alleles, but plasticity is conserved between *S. cerevisiae* and *S. paradoxus*, suggesting that there are compensatory *trans*-regulatory changes.

To estimate the proportion of divergence in expression level that is attributable to *cis*-acting changes, we considered the average log_2_ fold change in expression levels for all genes in all environments and asked how well expression differences between species were explained by expression differences between species-specific alleles in the F_1_ hybrids (Figure 8B). Using a regression analysis, we found that the coefficient of determination (R^2^) was 0.452, indicating that the relative allele-specific expression in F_1_ hybrids (i.e, *cis*-regulatory divergence) could account for nearly half of the differences in average expression level between species.

To estimate the proportion of divergence in expression plasticity that is attributable to *cis*-acting changes, we compared the correlation between species among timepoints in each environment to the correlation between species-specific alleles in the F_1_ hybrids over the same timepoints (Figure 8C). Correlation coefficients from all genes in all six environments were considered together for this analysis. We found that, apart from genes with plasticity that was conserved between species and alleles in F_1_ hybrids, the correlation between species-specific alleles in the F_1_ hybrid was a poor predictor of the correlation in expression among timepoints between species, resulting in an overall coefficient of determination (R^2^) of 0.124.

Based on these analyses of divergence in expression level (Figure 8B) and expression plasticity (Figure 8C), we infer that *cis*-regulatory changes explain much more of the divergence in expression level than the divergence in expression plasticity (R^2^ = 0.452 vs 0.124), suggesting that *trans*-regulatory divergence is primarily responsible for divergence in expression plasticity. (This finding is consistent with (Krieger et al. 2020), who reached the same conclusion using different methods of analysis.) Examples of this pattern are shown in Figures 8D and 8E, in which the expression level of each species and each allele in the F_1_ hybrids is shown for the 284 orthologs with divergent expression levels (Figure 8D) and the 289 orthologs with divergent expression plasticity (Figure 8E) in the diauxic shift environment. Note that the allele-specific expression of these genes in F_1_ hybrids was much more similar to the expression between species for expression level than expression plasticity (Figure 8D,E). Repeating these analyses for each of the six environments independently showed similar results (Figures S6 and S7), as did comparing expression plasticity using the scaled mean difference instead of the expression correlation among timepoints within an environment (Figure S7D). Although these data suggest that *cis*-regulatory divergence contributes more to divergence in expression level than expression plasticity, there are genes where divergence in expression plasticity was attributable primarily to *cis*-regulatory divergence (identifiable in Table S2 as genes that have both a low correlation between parental orthologs and between hybrid alleles). Such genes could potentially contribute to *trans*-regulatory divergence affecting expression plasticity of other genes.

To better understand the evolution of genes affected by both *cis*- and *trans*-regulatory divergence, we focused on cases where such changes have fully compensatory effects. That is, we examined genes where expression was conserved between species, but the species-specific alleles showed expression differences in the shared F_1_ hybrid cellular environment. To identify such cases, we considered all genes with a fold change less than 1.5 between species (log_2_ fold change = 0.585) as having conserved expression level. We then determined how many of these genes showed a fold change in expression level greater than 3 (log_2_ fold change = 1.585) between the species-specific alleles in the F_1_ hybrid. Depending on the environment, we found 41-85 genes with such fold changes in allele-specific expression in F_1_ hybrids (Figure 8F), indicating significant *cis*-regulatory divergence despite the conservation of this gene’s expression between species. This observation implies that there are also compensatory *trans*-regulatory differences between these two species that mask the effects of their *cis*-regulatory divergence.

To look for genes with compensatory *cis*- and *trans*-regulatory changes affecting expression plasticity, we assumed that genes with an expression correlation between species greater than 0.75 had conserved expression plasticity within an environment. We then defined genes with *cis*-regulatory divergence affecting expression plasticity as those with a hybrid expression correlation less than 0.5 (twice as far from 1 as our threshold). Depending on the environment, we found 72-223 genes that appeared to have compensatory *cis*- and *trans*-regulatory changes affecting expression plasticity (Figure 8G). In all six environments, we found more cases of compensatory *cis*- and *trans*-regulatory changes for expression plasticity than expression level using these thresholds (Figures 8F and G).

### Divergence of expression plasticity in the context of environment-specific regulatory networks inferred from data in Yeastract

To explore potential sources of *trans*-regulatory divergence contributing to divergence in expression plasticity as cells transition from one environment to another, we considered the regulatory connections between environment-specific regulators and their target genes based on experimental data assembled in Yeastract (Teixeira et al. 2018). Specifically, we downloaded regulatory matrices of transcription factors and their target genes for each environment, considering only regulatory connections for which there is (a) evidence of the regulator (transcription factor) binding near the potential target gene and (b) evidence that perturbation of the regulator affects expression of the potential target gene (either positively or negatively) in that environment. These data are shown as networks with edges connecting regulators and putative target genes for each of the six environments in Figure 9A. They are also summarized in Table S3, including the identities of each target gene and regulator.

**Figure 9.**
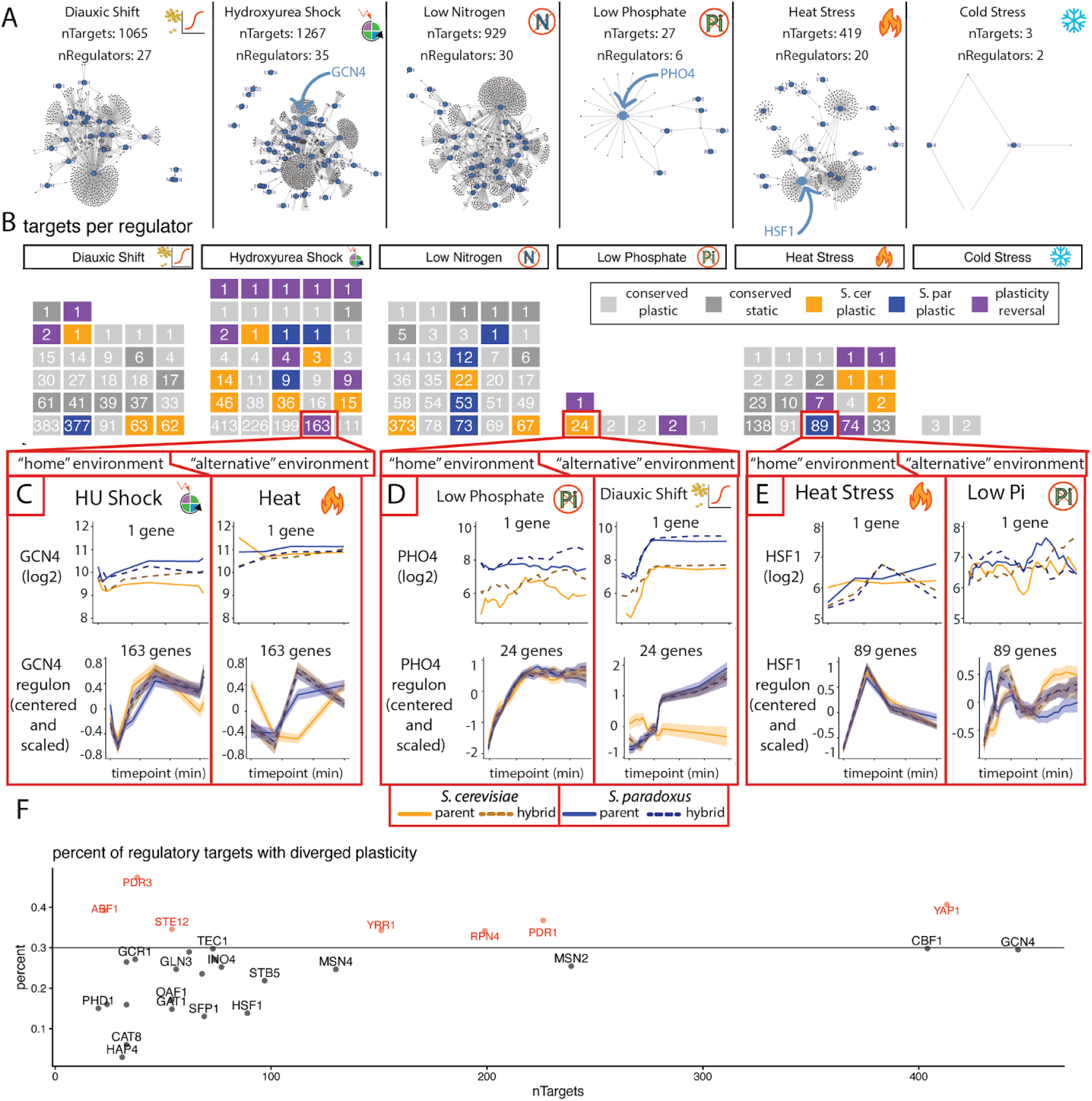
Expression profiles of target genes are often conserved despite divergent plasticity of key environment-specific regulators. A) Ball and stick models of environment-specific regulatory networks based on data downloaded from Yeastract for each of the six environments. Blue points indicate genes encoding transcription factors that function as regulators but can also themselves be targets of other regulators. Grey, smaller points indicate genes that are exclusively targets of regulation. The three transcription factors shown in panels C-E are labeled in these networks with arrows and larger labels: GCN4, PHO4, and HSF1. B) For each environment, the waffle plots show the number of target genes for each regulator (with at least one 1 target gene) in each environment-specific network. The color of each box indicates how expression plasticity of that regulator in that environment compares between species. Light grey indicates a regulator with conserved plasticity between species; dark grey indicates a regulator with conserved static expression; yellow indicates a regulator showing expression plasticity only in *S. cerevisiae;* blue indicates a regulator showing expression plasticity only in *S. paradoxus;* and purple indicates a regulator that showed a reversal in the direction of plasticity between species. C-E) Examples of three well-studied environment-specific regulators and their target genes (regulons) are shown: GCN4 (C), PHO4 (D), and HSF1 (E). In each case, in the top two plots, expression of the regulator in *S. cerevisiae* (yellow), *S. paradoxus* (blue), and allele-specific expression in F_1_ hybrids (dotted lines) are shown in the “home” environment (i.e., the environment where the regulator has a well-established role) and in an “alternative” (i.e., one of the other five environments). In the bottom two plots in each panel, average expression of the target genes for that regulator (as defined in the home environment) is shown for both the home and alternate environment for *S. cerevisiae* (yellow), *S. paradoxus* (blue), and allele-specific expression in F_1_ hybrids (dotted lines). Shading around each line represents the 95% confidence interval of the mean. For GCN4, PHO4, and HSF1, respectively, these target gene sets included 163, 24, and 89 genes. F) The percentage of genes in each regulon with divergent plasticity in the home environment is shown for the 41 regulators with at least 20 target genes. Each point is one environment-specific regulator from the Yeastract database. The percentage of the regulon with divergent plasticity in the home environment (y-axis) is plotted by the number of genes in the regulon of each transcription factor (x-axis). The horizontal line shows the 30% cutoff value used in the text.

The number of environment-specific regulators in this dataset ranged from 2 to 56, and the number of target genes ranged from 3 to 1270, resulting in environment-specific networks of varying complexity (Figure 9A). These differences in network complexity might reflect differences in data availability or curation, as well as differences in yeast biology, among environments. Note that these networks are based solely on data collected in *S. cerevisiae*, so they do not include any regulatory relationships unique to *S. paradoxus*. Despite these limitations, we used these networks to explore how divergent expression of a transcriptional regulator might impact expression of downstream genes.

Among all six environments, Yeastract identified 120 environment-specific regulators with at least one putative target gene identified in our dataset (Table S4). This set includes 94 unique transcription factors, as some of these regulators were identified in multiple environments. For example, GCN4 has documented regulatory interactions in the diauxic shift, hydroxyurea shock, and low nitrogen environments. Focusing on expression plasticity, we found that 42 of the 120 (35%) environment-specific regulators showed divergence in expression plasticity between species (Figure 9B). 16 of these 42 regulators showed changes in expression as cells transitioned to a new environment in *S. cerevisiae* but not *S. paradoxus*, and 9 showed the opposite pattern. The remaining 17 regulators showed a reversal of plasticity between species, with expression increasing during acclimation to a new environment in one species and decreasing in the other (Figure 9B).

To determine how these changes in expression of regulatory proteins relate to expression of their target genes, we compared expression of the regulator over experimental time in each environment to the expression of the set of genes predicted by the data in Yeastract to be direct targets of that regulator. This analysis was motivated by the expectation that divergent expression of an important environment-specific regulator might cause similar expression divergence in the group of genes regulated by that transcription factor (i.e., its “regulon”). Focusing on the 14 transcription factors that have both divergent plasticity in their “home” environment (i.e., the environment in which they were identified as a regulator in the regulatory matrix) and more than 20 predicted target genes in the same environment (Figure 9B, highlighted in Table S3), we found that this was not generally the case. Rather, for 6 of these 14 transcription factors, expression of genes in their regulon was more highly correlated between species in their “home” environment than in any of the other five environments tested (highlighted in green in Table S3). These cases included PHO4 in the low phosphate environment (Figure 9D) and HSF1 in the heat stress environment (Figure 9E). An additional 6 transcription factors showed a correlation in expression between species in their home environment equal to or greater than the average expression correlation among all six environments (highlighted in yellow in Table S3), including GCN4 in the hydroxyurea shock environment (Figure 9C). Our results here are consistent with recent findings that changes in expression of regulators do not always affect the expression of their downstream targets (Domingo et al. 2026).

Finding that divergent plasticity of an environment-specific regulator is a poor predictor of divergent plasticity in its target genes could suggest that other changes between species in the regulatory network have evolved to compensate for the divergence of these environment-specific regulators. Such compensation would presumably evolve under stabilizing selection. However, this conserved plasticity of target genes could also be explained by the conservation of other regulators that combinatorially control expression of these target genes. Indeed, we found that in five of the six environments, genes with conserved expression plasticity tended to have more interactions in the environment-specific regulatory network than genes with divergent expression plasticity (Fisher’s Exact Test p-values < 1×10^-5^, Figure S8). The one environment where this relationship was not seen was the cold stress environment, which contained only five documented regulatory interactions and thus had little power to detect this relationship.

Ignoring whether the regulator itself showed evidence of plasticity divergence, we also examined the conservation and divergence of expression plasticity for regulators with sets of at least 20 target genes in each environment. We found that 27 (66%) of the 41 regulators in environment-specific networks with at least 20 genes in their regulons (Table S4) showed greater conservation of environment-specific expression plasticity than expected by chance based on permutation testing (Figure S9). For this analysis, a permutation test was run for each of these 41 environment-specific regulators by randomly sampling the same number of genes as in the regulator’s environment-specific regulon, recording the percentage of these random genes with divergent plasticity in the regulator’s home environment, and repeating this process 1000 times to form a null distribution. We then compared the observed proportion of target genes with divergent plasticity of the regulator in its home environment to the corresponding null distribution and considered the regulon to be enriched for genes with conserved plasticity if the observed value was less than 95% of the values in the null distribution (Figure S9).

To search for regulators with target genes showing divergent expression plasticity most often between *S. cerevisiae* and *S. paradoxus*, we focused on the 29 unique transcription factors that appeared among the list of 41 environment-specific regulators with at least 20 target genes in at least one environment (Table S4). Of these 29, we found that 7 had at least 30% of their target genes with divergent expression plasticity (Figure 9F, red points and labels). These transcription factors were ABF1, STE12, YAP1, RPN4, PDR1, PDR3, and YRR1. Interestingly, four of these seven transcription factors (YAP1, RPN4, PDR1, and PDR3) have been identified as key regulators that allow *S. cerevisiae* to metabolize the toxic compound 5-Hydroxymethylfurfural (HMF) (Ma and Liu 2010), and a fifth, YRR1, has been implicated in resistance to another toxic compound, 4-nitroquinoline 1-oxide (4NQO) (Gallagher et al. 2014) and recognizes a DNA binding site that is similar to that of PDR1 and PDR3 (Le Crom et al. 2002). The target genes for the above five PDR regulators (YAP1, RPN4, PDR1, PDR3, and YRR1), as well as the subset of these target genes with divergent expression plasticity, have minimal overlap among regulators (Figure S10).

Interestingly, all five of these transcription factors showed evidence of divergent expression level rather than divergent expression plasticity. In all cases, higher expression was seen in *S. paradoxus* than in *S. cerevisiae,* as well as for the *S. paradoxus* allele in F_1_ hybrids (Supplemental Table 4), indicating that this expression divergence is most likely caused by *cis*-regulatory divergence in each of these transcription factor genes. Whether and how these changes in transcription factor expression level might contribute to divergence in environment-specific expression plasticity of their target genes remains unknown, but this observation warrants further investigation. Regardless of molecular mechanisms that might ultimately be discovered, the divergence in expression level and expression plasticity seen for these transcription factors and their targets suggests that *S. cerevisiae* and *S. paradoxus* might have evolved differences in their response to drugs that trigger the pleiotropic drug resistance network. Indeed, one of these genes, YRR1, has previously been found to harbor genetic variation in its coding and promoter sequences that contributes to variation in gene expression and growth rate of *S. cerevisiae* strains (Gallagher et al. 2014), suggesting that a similar mechanism might contribute to phenotypic divergence between *S. cerevisiae* and *S. paradoxus*.

## Conclusion

The regulation of gene expression in response to the environment is a fundamental property of all organisms, and evolutionary changes in this regulation can help explain phenotypic differences among species. A prior study examining the evolution of gene expression between *S. cerevisiae* and *S. paradoxus* found that expression level and expression plasticity are evolving independently, with effects of genetic changes altering gene expression level often acting in *cis* and causing similar differences in expression level among environments (Krieger et al. 2020). Genetic changes affecting expression plasticity were found to be more often in *trans* and have more variable effects among environments (Krieger et al. 2020). Here, we combined the data from (Krieger et al. 2020) with data from (Fay et al. 2023) and used it to investigate how *S. cerevisiae* and *S. paradoxus* respond transcriptionally as they acclimate to each of six different environments. We found that divergence in expression level seen consistently among timepoints following a transition to one new environment was generally consistent for the same gene in all six environments. This finding contrasts with divergence in expression plasticity (defined here as differences among timepoints after moving to a new environment), which was most often observed only in one of the six environments for a given gene. The different contributions of *cis*- and *trans*-regulatory changes to divergence in expression plasticity and expression level (which we also saw among timepoints within each environment) likely explain this difference in context-dependency of these two types of regulatory divergence.

Among genes showing divergence in expression plasticity, we identified cases where only one species increased or decreased expression following the transition to a new environment, suggesting that this gene has either acquired a derived response in one species or lost a pre-existing response in the other. We are unable to distinguish between these possibilities because we only analyzed the two species; we did not also examine an outgroup. In other cases, we found that genes increased expression in response to a new environment in one species but decreased expression in response to the same environment in the other species. These reversals in plasticity might involve an evolutionary switch of regulators from activation to repression. We found that divergence in the expression plasticity of transcription factors regulating sets of genes within a particular environment generally did not translate into similar changes in expression of those downstream genes. Rather, expression of genes regulated by a transcription factor with divergent plasticity in a particular environment tended to remain similarly expressed between species. Despite most environment-specific regulons having more genes with conserved expression plasticity than expected by chance, we found that five regulators in the pleiotropic drug resistance network all had *cis*-regulatory divergence leading to higher expression levels in *S. paradoxus* than *S. cerevisiae* as well as a larger percentage of target genes with divergent expression plasticity than other regulators, but it remains to be seen whether there is a causal relationship between these observations and whether these changes contribute to phenotypic divergence.

The environment-specific divergence in regulatory networks between *S. cerevisiae* and *S. paradoxus* uncovered in this study provides a foundation for future studies investigating the specific molecular and genetic changes responsible for these evolutionary changes in expression plasticity. Past work identifying specific *cis*- and *trans*-regulatory changes responsible for such changes in yeast gene regulatory networks has used much more distantly related species (Tsong et al. 2006; Baker et al. 2011; Britton et al. 2020), and tracking down the specific mechanisms responsible for this divergence between *S. cerevisiae* and *S. paradoxus* over a much shorter evolutionary timescale will provide important complementary insight that contributes to a more comprehensive understanding of evolutionary processes.

## Methods

### RNA-seq datasets and reference genome sequences

RNA-seq data analyzed in this study comes from (Krieger et al. 2020) and (Fay et al. 2023). Raw RNA-seq reads from Krieger et al. (2020) were downloaded from the NCBI BioProject database (https://www.ncbi.nlm.nih.gov/bioproject/) using accession number PRJNA592756. Raw RNA-seq reads from Fay et al. 2023 were downloaded from the same database using accession number PRJNA909640. From the (Krieger et al. 2020) dataset, only samples from wild-type genotypes were included in our analysis; mutant genotypes also analyzed in this study were excluded. From the (Fay et al. 2023) dataset, only reads from *S. cerevisiae* and *S. paradoxus* were included in our analysis; reads from the other two species included in this study (*S. kudriavzevii* and *S. uvarum*) were excluded. Because (Fay et al. 2023) collected their RNA-seq data from four different species in a common pool, excluding these reads required first mapping them to a concatenated genome that included all four species. Genome assemblies and gene annotations for *S. cerevisiae* (S288C) and *S. paradoxus* (CBS432) used to map RNA-seq reads were downloaded from https://yjx1217.github.io/Yeast_PacBio_2016/data/ (Yue et al. 2017). Genome assemblies for *S. kudriavzevii* (GCA 947243775.1) and *S. uvarum* (GCA 027557585.1) from (Langdon et al. 2019) were downloaded from NCBI.

9 of 705 RNA-seq samples with less than 100,000 total reads were excluded from analysis. Seven of these samples came from the hydroxyurea shock environment (3 *S. cerevisiae*, 2 *S. paradoxus*, and 2 hybrid samples at various timepoints). We also excluded 2 hybrid samples from the hydroxyurea shock environment because they had substantially different library sizes for the *S. cerevisiae* and *S. paradoxus* (Spar) alleles (1,665,985 Scer reads vs. 19,058 Spar reads and 425,558 Scer reads vs. 190,710 Spar reads), suggesting that not all cells sampled were F_1_ hybrids.

### Mapping RNA-seq reads

We started with the raw RNA-seq reads rather than processed gene counts from (Krieger et al. 2020) and (Fay et al. 2023) to ensure that read mapping was comparable for all data analyzed in this study. But there are two other differences between the two studies to note: (a) they used different strains of *S. cerevisiae* (S288c vs. T73) and *S. paradoxus* (CBS432 vs. N17), and (b) they used different strategies for making cDNA libraries for RNA-seq (3’ Tag-Seq vs. full transcript sampling). To account for the latter difference, read counts from (Fay et al. 2023) were normalized for gene length, whereas read counts from (Krieger et al. 2020) were not (because each read comes from a different mRNA transcript).

STAR v2.7.11b (Dobin et al. 2013) was used to both align and quantify RNA-seq reads. Prior to aligning reads, we generated reference genome indices using STAR in *–runMode genomeGenerate* with the following parameters: *–genomeSAindexNbases 11* (adjusts the index complexity for the 12MB yeast genome size), *–sjdbGTFfeatureExon gene* (treats the full gene as an exon; introns are rare in *S. cerevisiae*), and *–sjdbGTFtagExonParentGene Name* (use gene names as the primary ID). When analyzing the 75 bp reads from (Fay et al. 2023), we also included the parameter *–sjdbOverhang 74,* which sets the search window for splice junctions, adjusting sensitivity for differences in read length between the two studies.

The 50 bp 3’ Tag-Seq reads from (Krieger et al. 2020) were mapped to a concatenated genome containing *S. cerevisiae* and *S. paradoxus* using STAR with the following parameters:

*–alignEndsType EndToEnd* (requires mapping of the full 50 bp read),

*–outFilterScoreMinOverLread 0* (turns off default score thresholds),

*–outFilterMatchNminOverLread 0* (turns off default mismatch threshold of 10),

*–outFilterMismatchNmax 4* (sets mismatch threshold of 4/50 bp, or 8%), *–scoreGap -10* (strongly penalizes gaps in alignments). STAR was run in *–quantMode GeneCounts* mode to concurrently count reads within 500 base pairs of the transcription end site (±500 TES). On average, ∼60% of reads (∼1 million reads) mapped uniquely to one genome or the other in each sample. Only uniquely mapped reads overlapping ±500 of an annotated transcription end site (TES) were retained.

The 75 bp RNA-seq reads from Fay et al. 2023 were mapped to a concatenated genome including *S. cerevisiae*, *S. paradoxus*, *S. kudriavzevii,* and *S. uvarum* using STAR with the following parameters: *–alignEndsType EndToEnd* (requires mapping of the full 75 bp read),

*–outFilterScoreMinOverLread 0* (turns off default score thresholds),

*–outFilterMatchNminOverLread 0* (turns off default mismatch threshold of 10),

*–outFilterMismatchNmax 6* (sets mismatch threshold of 6/75 bp, or 8%), *–scoreGap -10* (strongly penalizes gaps in alignments). STAR was run in *–quantMode GeneCounts* mode to concurrently count reads overlapping an annotated gene. While these reads were from paired-end data, they were mapped as single-end for consistency with the (Krieger et al. 2020) data. However, because read 1 and read 2 had highly correlated gene counts (average R=0.98, minimum R=0.9), we used the average count for each set of paired end reads. An average of 84% of reads (∼10 million reads) mapped uniquely in each sample. Only uniquely mapped reads overlapping an annotated gene were retained.

### Assessing mapping bias

We assessed allele-specific mapping bias by measuring the percent of reads collected from *S. cerevisiae* that mapped to the *S. paradoxus* and vice versa. This was only possible to assess with the (Krieger et al. 2020) data because (Fay et al. 2023) sequenced their parental DNA samples from both species in a single pool. To assess mapping bias on a gene-by-gene basis, we counted the total number of reads that mapped to each ortholog in each parental sample. We found that 36 of 4491 genes in *S. cerevisiae*, 50 of 4491 genes in *S. paradoxus*, and 3 of 4491 genes in both species (TDH1, ESC2, and YCL048W-A) had less than 90% of reads mapping to the correct parental allele. These genes were labeled as “biased” in the level column and excluded from analysis (but retained in Table S1 for informative purposes). Note that we included the *–alignEndsType EndToEnd* parameter, preventing soft-clipping in our mapping protocol, because when we allowed STAR to use its default soft-clipping parameter, over 500 genes showed significant mapping bias, indicating that the soft-clipping was causing many reads to map to the wrong species’ allele.

### Normalizing count data

Sequencing read count data was normalized to counts per million. For the 3’ Tag-Seq data from (Krieger et al. 2020), which is stranded and polyA-selected, one read is equivalent to one transcript, so normalizing each gene’s expression in each sample was done as follows:

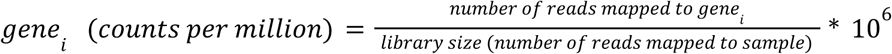

The (Fay et al. 2023) RNA-seq data was paired-end, stranded, and polyA-selected, with reads expected to be distributed over the length of each transcript. Consequently, to convert the number of sequence reads mapping to a gene to a comparable counts per million metric, we normalized for transcript length using the following equation from (Zhao et al. 2020):

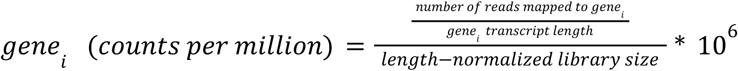

where

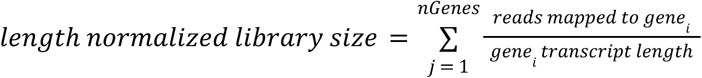

Finally, we excluded genes with fewer than 30 counts per million in any experiment or for either allele in F_1_ hybrids. This filter reduced the number of genes analyzed from 5359 to 4997.

### Generalized linear model to determine divergence in level within an environment

For each gene *i*, in each experiment *j*, we fit the following negative binomial generalized linear model with a log link function using the *glm.nb* function from the R package *MASS* (Venables and Ripley 2002):

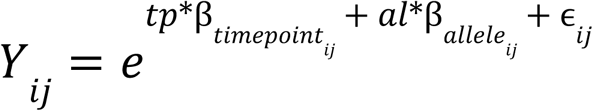

Where *Y_ij_* is the read count (in counts per million) of gene *i* in experiment *j* at a given timepoint for a given allele (*S. cerevisiae* or *S. paradoxus*), *ɛ_ij_* is the response residual (actual – predicted value of *Y_ij_*), and 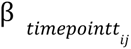 and 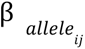 are the coefficients for the continuous variable timepoint (*tp*, ranging from 0 to around 1000 depending on the experiment, in minutes) and the categorical variable allele (*al*, 0 for *S. paradoxus*, 1 for *S. cerevisiae*).

From this model, we extracted each β_ij_, the estimated (natural) log-fold change associated with switching from the *S. paradoxus* to the *S. cerevisiae* allele at any given timepoint for gene *i* in experiment *j*. We converted this to a log_2_ fold change,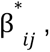 to be consistent with prior studies using the change of base formula:

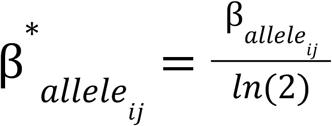

A gene *i* in experiment *j* was determined to be diverging in expression level if 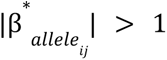 and its associated Wald test P-value was less than 0.05 after adjusting P-values for multiple testing using a false discovery rate according to the Benjamini-Hochberg procedure (Benjamini and Hochberg 1995).

The proportion of genes identified as diverging in expression level was highly dependent on the chosen effect size threshold—without an effect-size threshold, roughly 3 times as many genes were diverging in expression level versus expression plasticity. Conversely, with a stronger threshold (i.e., requiring that a gene be expressed twice as high in one species to be considered diverging in level), roughly 10 times more genes were diverging in plasticity than level (Figure S3). Significance of the Wald test and a fold change threshold of 1.5 were used for identifying genes with divergent expression levels in the main text, except where noted.

### Correlation clustering to identify divergence in plasticity

Hierarchical correlation clustering was carried out using custom R code inspired by the R package Weighted Gene Co-expression Network Analysis (WGCNA, (Langfelder and Horvath 2008)). The input data for correlation clustering is a counts table where rows are genes, columns are samples, and all values are in counts per million (Cleaned_Counts.RData on GitHub). We performed the clustering separately for samples from each of the six experiments (Diauxic Shift, Hydroxyurea Shock, Low Nitrogen, Low Phosphate, Heat Stress, and Cold Stress), using each gene’s mean normalized read count for all replicates at each timepoint. The timepoints taken in the low phosphate and diauxic shift experiments did not have replicates, so we used a moving average instead of the mean among replicates, making each timepoint the average of itself and its two neighboring upper and lower timepoints (i.e., a sliding window of 5).

For each of the six experiments, prior to clustering, we first identified two groups of genes: 1) lowly expressed genes whose average expression was less than 30 counts per million, and 2) lowly varying genes whose average expression was greater than 30 counts per million but whose dispersion (var/mean) was less than 1. We determined these two thresholds by plotting each gene’s log_2_(variance(expression)) against its log_2_(mean(expression)). We set thresholds on this plot to separate out as many genes as possible with abnormally high variance given their mean expression (which tended to be lowly expressed genes) and genes with abnormally low variance given their mean (lowly varying). These lowly-varying genes are labeled as “static” in Table S1. Note that as described in the data cleaning section, we had already filtered out genes with less than 30 counts per million in both species in all six experiments. This step removed additional genes that were lowly expressed in the focal experiment in one (or both) species. The lowly expressed genes were omitted in Table S1. Both lowly varying and lowly expressed genes were excluded from the hierarchical clustering for that environment.

Using the remaining genes, we constructed a single correlation matrix for each of the six experiments. Each *[i,j]* entry in the correlation matrix was the expression correlation of gene *i* and gene *j* in that experiment, where *i* and *j* can be orthologs of the same gene—one from *S. cerevisiae* and one from *S. paradoxus*. To assign these genes to clusters with correlated expression, we performed bottom-up hierarchical clustering on the additive inverse of the correlation matrix (i.e., a correlation matrix where every value was multiplied by -1, because hierarchical clustering attempts to minimize distance between objects in clusters).

In bottom-up hierarchical clustering, each gene begins as its own cluster, and at each step the pair with the strongest correlation is joined into the same cluster. After the first step, groups of genes that are joined are summarized by the average of their pairwise correlations with a candidate next gene or gene group to determine whether they should be the next pair to be joined. Once all genes used for clustering in an environment were joined into one gene tree, we cut the tree to determine the number of clusters. Hierarchical clustering does not produce a specified number of clusters, unlike k-means clustering. Rather, we cut our gene tree three times by cutting our gene trees at the top, second-to-top, and third-to-top branch, creating sets of genes with 2, 3, or 4 clusters (in addition to the static cluster), respectively. For each environment, the clustering results were bootstrapped by clustering a randomly selected 75% of genes 100 times and taking the most frequent cluster assignment of each gene as its cluster.

### Yeastract regulatory matrices

Binary regulatory matrices were downloaded from <u>Yeastract.com</u> (Teixeira et al. 2018) for each of the six environments by selecting the following environmental conditions from the dropdown list options available in Yeastract: (a) Diauxic Shift: Carbon source quality/availability > non-fermentable carbon source. There is also a “Diauxic Shift” environment available in Yeastract, but it only includes 30 regulatory connections. (b) Hydroxyurea Shock: Stress > Drug/chemical stress exposure, (c) Low Nitrogen: Nitrogen source quality/availability > Nitrogen starvation/limitation, (d) Low Phosphate: Ion/metal/phosphate/sulfur/vitamin availability > Phosphate limitation, (e) Heat Stress: Stress > Heat shock, and (f) Cold Stress: Stress > Cold shock. We also downloaded a reference regulatory network by selecting the “Unstressed log-phase growth” environment (no subgroup). Regulatory connections in these networks required evidence of both “DNA binding” (e.g., ChIP-seq data showing evidence of a TF binding near a target gene) and “expression” (e.g., the target gene changes expression when the TF is deleted). We used all *S. cerevisiae* transcription factors in Yeastract (with the “Consider S. cerevisiae TF list” option) and entered a list of potential target genes for these TFs (every gene that appears in Supplementary Table 1 at least once), in which genes identified as lowly expressed in all environments were excluded. Networks shown in Figure 9A were produced using the *igraph* package in R (Csardi and Nepusz 2006).

## Code Availability

All code used in these analyses is available for viewing at the GitHub Pages site (<u>wyldtype.github.io/Plasticity/</u>) and for download at the GitHub Repository (<u>github.com/wyldtype/plasticity</u>). The code in this project used the R packages WGCNA (Langfelder and Horvath 2008) and MASS (Venables and Ripley 2002) for data analysis and ComplexHeatmap (Gu et al. 2016) and circlize (Gu et al. 2014) for data visualization.

## Acknowledgements

We are grateful to Naama Barkai (Weizmann Institute of Science) for sharing data before publication and for many insightful and collaborative discussions over the last six years. Mo Siddiq for his thoughts on how to communicate our findings and Luis Zaman for contributing his expertise in biological network theory. We also thank all members of the Wittkopp lab for valuable discussions and comments on drafts of this manuscript. Funding for this work was provided by awards from the National Science Foundation (MCB-1929737) and the Michigan Israel Partnership Award (G024016-336246) to PJW as well as the National Institutes of Health Training Grant (T32HG000040) support to ACR.

## Notes

### Competing Interest Statement

The authors have declared no competing interest.

