## Supplementary Figures for "Evolution of regulatory networks controlling plasticity in gene expression between *Saccharomyces cerevisiae and Saccharomyces paradoxus*"

by

Anna Redhuis, Naama Barkai, and Patricia J. Wittkopp

Contents:

**Figure S1.** Inference of divergent expression plasticity with 2, 3, or 4 clusters per environment

**Figure S2.** Divergence in expression level and plasticity for all 6 environments

**Figure S3.** Impact of changing the threshold for calling divergent expression level on the proportion of genes in expression divergence categories

**Figure S4.** Impact of changing the threshold for calling divergent plasticity on the proportion of genes in expression divergence categories

**Figure S5.** Genes with divergent plasticity in more than one environment

**Figure S6.** Divergence in expression level is largely recapitulated in the $F_1$ hybrid allele-specific expression

**Figure S7.** Divergence in expression plasticity is rarely recapitulated in the $F_1$ hybrid allele-specific expression

**Figure S8.** Genes with conserved plasticity tend to have more regulatory connections in environment-specific networks

**Figure S9.** Conservation of expression plasticity in environment-specific regulons with at least 20 genes

**Figure S10**. Minimal overlap of genes with divergent expression plasticity that are targets of five transcription factors involved in the pleiotropic drug response network

**Table S1.** Average expression levels and expression plasticity between *S. cerevisiae* and *S. paradoxus* in six environments

**Table S2.** Expression differences between species and between alleles in $F_1$ hybrids in six environments

**Table S3.** Summary of environment-specific regulatory interactions between transcription factors and target genes from Yeastract

**Table S4.** Expression divergence of environment-specific transcriptional regulators and their regulons

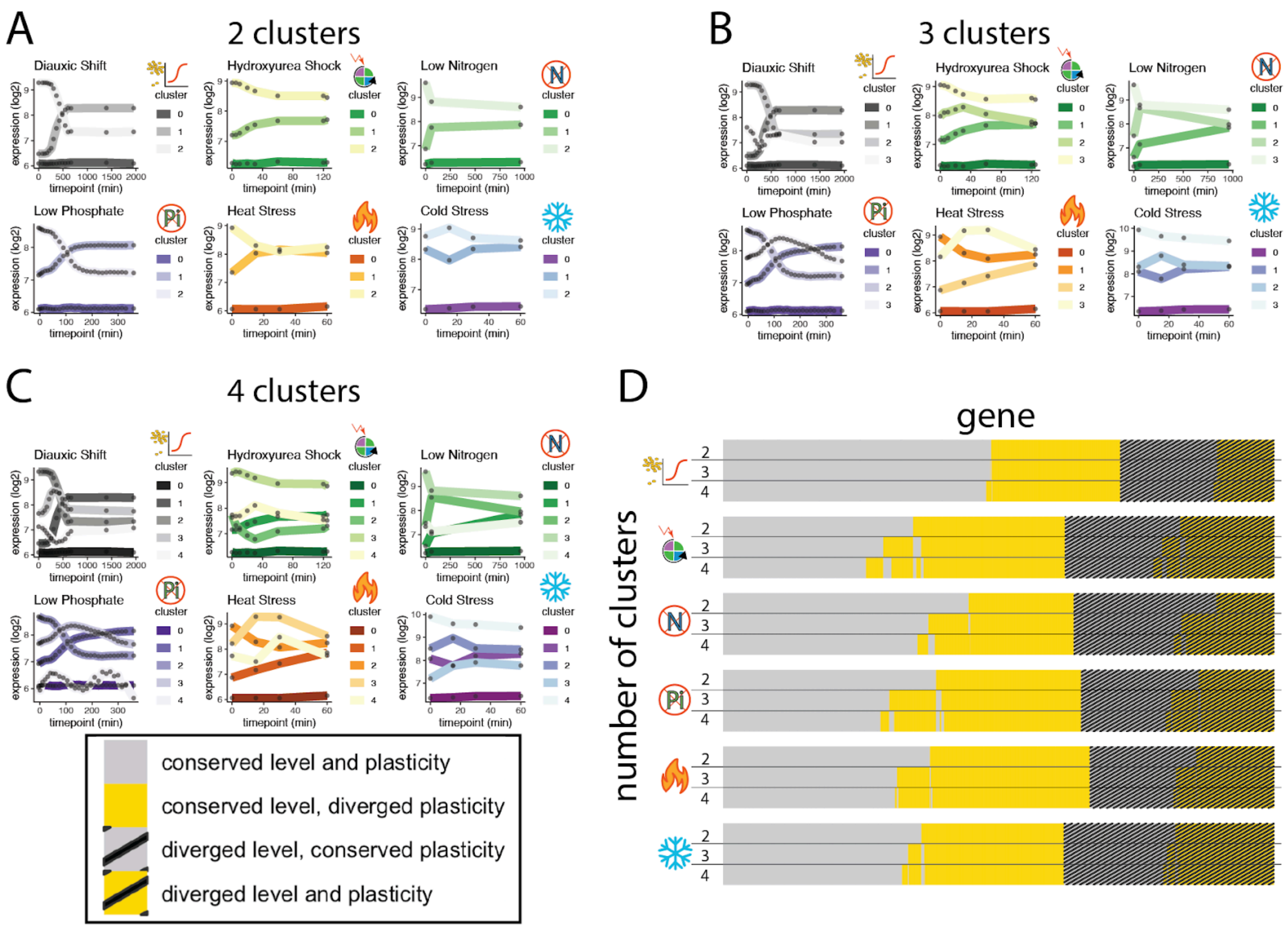


**Figure S1**. **Inference of divergent expression plasticity with 2, 3, or 4 clusters per environment.** (A) This panel is a reproduction of the data shown in Figure 4, provided here for comparison with panels B and C. In each panel, the lines show the average expression of all genes in each cluster, with black dots on the lines showing the timepoints at which samples were taken. Cluster 0 is defined as static, cluster 1 is defined as increasing, and cluster 3 is defined as decreasing. (B) The same analysis shown in A, but after clipping the hierarchical clustering tree at a level that divided genes into 3 expression plasticity clusters in each environment rather than 2. Cluster 0 is most similar to the static cluster in A, but clusters 1, 2, and 3 show more complex patterns than clusters 1 and 2 in A. (C) The same analysis shown in A, but after clipping the hierarchical clustering tree at a level that divided genes into 4 expression plasticity clusters in each environment. Cluster 0 is most similar to the static cluster in A, but clusters 1, 2, 3, and 4 show more complex patterns than clusters 1 and 2 in A. (D) The relative proportions of genes defined as having conserved or divergent expression levels and expression plasticity for all three clustering scenarios are shown for all six environments. For each environment, the genes are shown in the same order from right to left for the analyses using 2, 3, or 4 clusters. The similarity of these plots for 2, 3, and 4 clusters in the same environment suggests that the decision to analyze these data using only 2 expression plasticity clusters (increasing and decreasing) had little effect on the interpretation of whether a gene had divergent expression level and/or divergent expression plasticity.  Logos represent the six different environments, as defined in Figure 2.

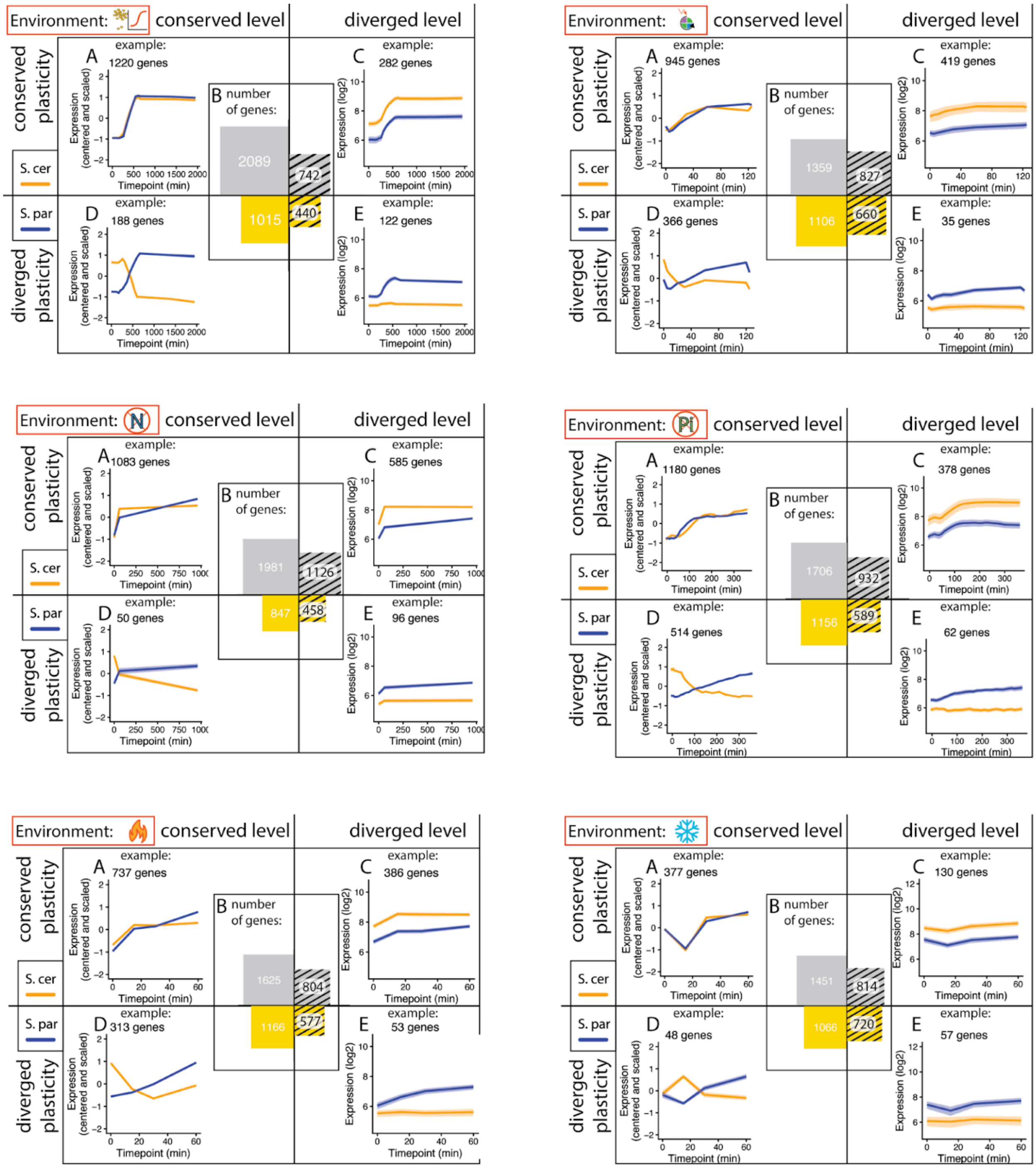


**Figure S2**. **Divergence in expression level and plasticity for all 6 environments.** Results from each of the six environments are shown that are comparable to Figure 5 for the diauxic shift environment. Figure 5 is reproduced here for comparison as the top left panel. The same analyses are shown for the other 5 environments in the other five panels, with the logos (as defined in Figure 2) identifying each environment. In all 6 panels, subpanel A shows the number of genes in each of the four conserved or diverged categories for expression level and plasticity. Subpanel B shows the average expression level of genes with conserved expression level and plasticity that increased expression in both species. Subpanel C shows the average expression of genes that have a higher expression level in *S. cerevisiae* and increased expression at the diauxic shift in both species. Subpanel D shows the average expression of genes that have conserved expression level, decrease expression in *S. cerevisiae,* and increase expression in *S. paradoxus*. Subpanel E shows the average expression of genes with a higher expression level in *S. paradoxus* and an increase in expression at the diauxic shift only in *S. paradoxus*. In subpanels B-D, shading around the lines are error ribbons that indicate 95% confidence intervals for the mean and the number of genes included in the line plot is shown.

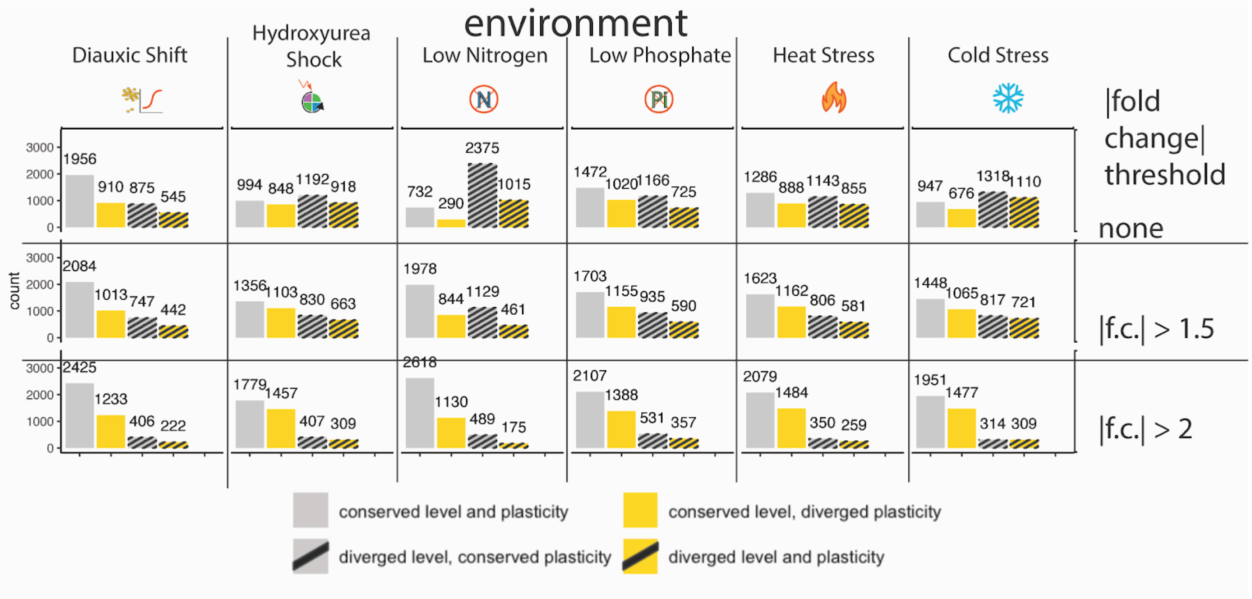


**Figure S3. Impact of changing the threshold for calling divergent expression level on the proportion of genes in expression divergence categories.** In each of the 6 environments, the proportions of genes belonging to each class of expression divergence are shown based on analyses using different thresholds to classify genes as showing divergent expression level. Top row: any gene with a Wald test adjusted P-value < 0.05 is considered to be diverging in expression level. Middle row: genes with a Wald test adjusted P-value < 0.05 and an estimated fold change in mean expression at least 1.5x higher in one species than the other are considered to be diverging in expression level. Bottom row: genes with a Wald test adjusted P-value < 0.05 and estimated fold change in mean expression at least 2x higher in one species than the other are considered to be diverging in expression level. Numbers show the number of genes in each category in each environment with each threshold for calling divergence in expression level.

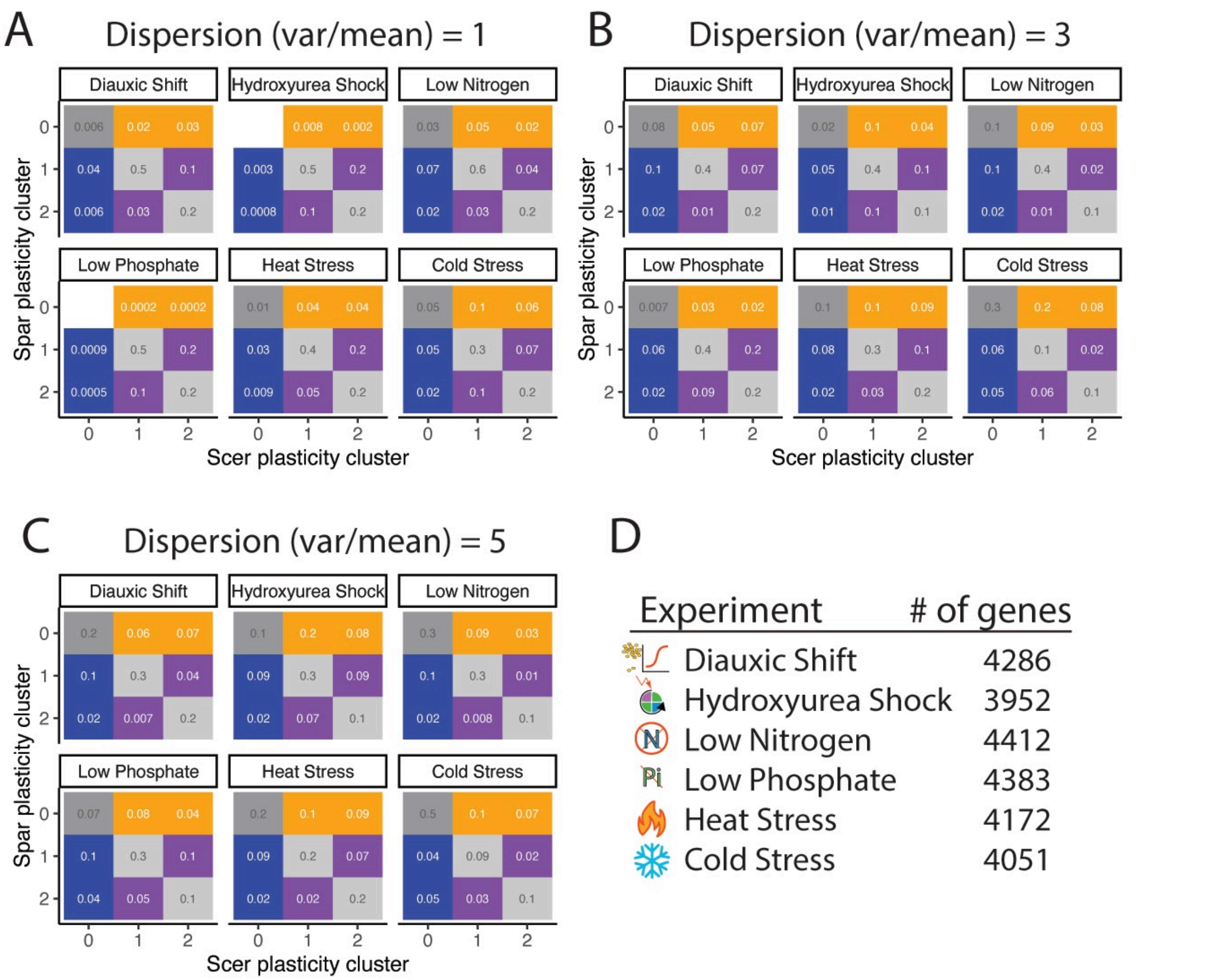


**Figure S4. Impact of changing the threshold for calling divergent plasticity on the proportion of genes in expression divergence categories.** (A-C) For each of the six environments, the proportion of genes with orthologs in each combination of plasticity clusters is shown. Cluster 0 = conserved plasticity; cluster 1 = increasing in expression across timepoints; and cluster 2 = decreasing in expression across timepoints. Grey boxes indicate genes with expression plasticity conserved between species; yellow boxes indicate genes that showed plasticity only in *S. cerevisiae*; blue boxes indicate genes that showed plasticity only in *S. paradoxus*; and purple boxes indicate genes that showed a reversal in plasticity between species, with one species ortholog showing expression increasing among timepoints in an environment and the other species showing expression decreasing among timepoints in the same environment. (A) Proportion of genes in each category when a threshold of variance/mean = 1 was used to decide which genes showed static versus plastic expression. (B) The same analysis as in (A) but with a variance/mean =3 threshold; these data are repeated from Figure 6D. (C) The same analysis as in (A) but with a variance/mean = 5 threshold. Note that the proportion of genes considered to have static (non-plastic) expression went up as this variance/mean threshold was raised, making the data shown in panel C the most conservative for identifying genes with expression plasticity. (D) The number of orthologous gene pairs analyzed in each environment.

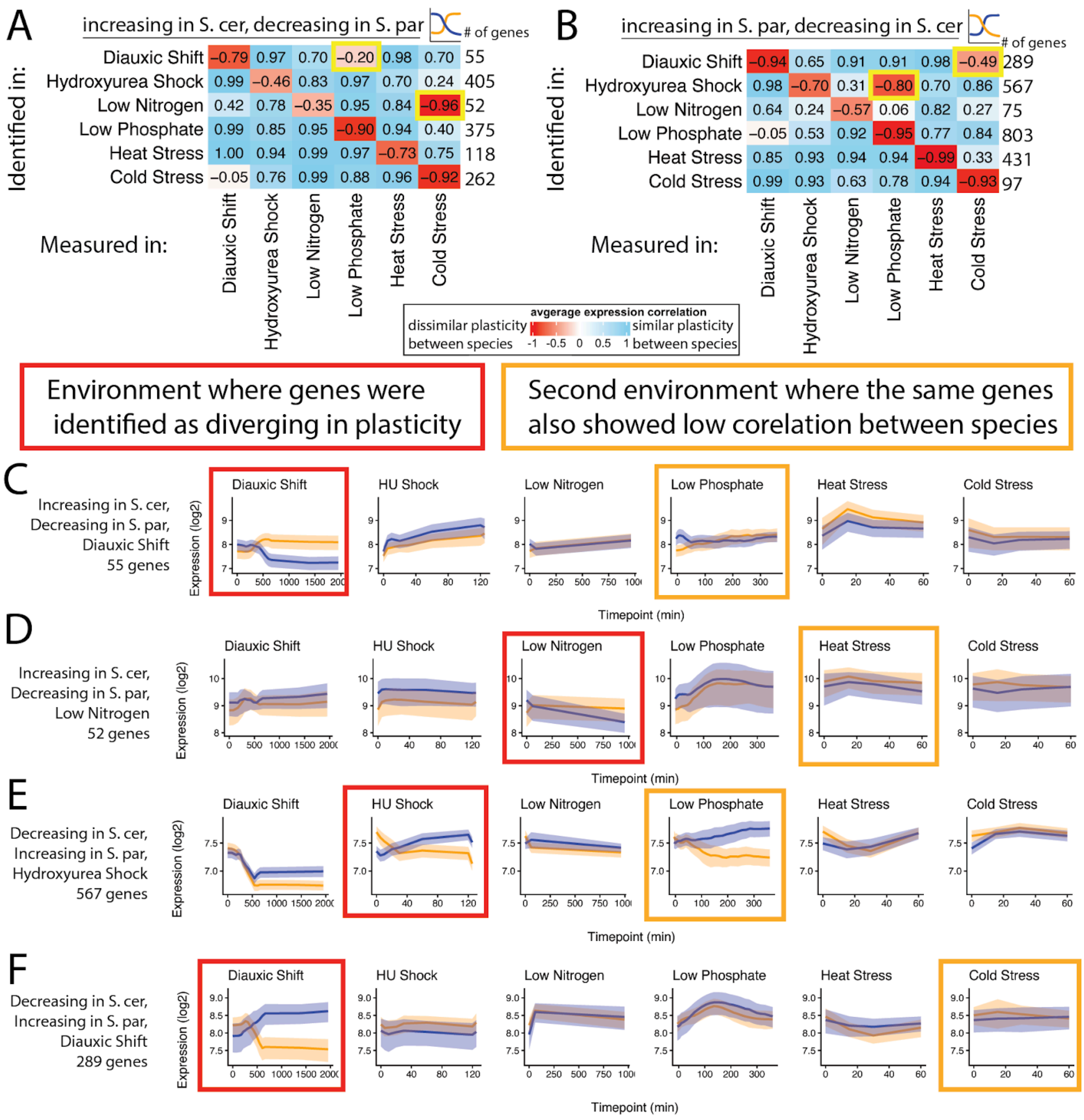


**Figure S5. Genes with divergent plasticity in more than one environment.**

(A) The matrix and color coding are reproduced from Figure 7A, top panel, showing the correlation in expression across timepoints within an environment between *S. cerevisiae* and *S. paradoxus* for genes classified as increasing over time in *S. cerevisiae* and decreasing in *S. paradoxus*. For each cell, the gene set considered was defined by showing a reversal in plasticity in the environment shown to the left of the matrix (i.e., “Identified in”) and then analyzed using data from the environments shown at the bottom of the matrix (“Measured in”). Two cells are highlighted with a yellow border because they showed a negative correlation with a correlation coefficient less than -0.1 in the “Measured in” environment. One of these cells includes a set of genes identified in the diauxic shift environment and measured in the low phosphate environment (shown in more detail in panel C) and one includes a set of genes identified in the low nitrogen environment and measured in the cold stress environment (shown in more detail in panel D). (B) Same as panel A except that the genes with plasticity reversals considered had expression increasing in *S. paradoxus* and decreasing in *S. cerevisiae.* The matrix and color coding are reproduced from Figure 7A, bottom panel. Again, two cells are highlighted with a yellow border because they showed a negative correlation with a correlation coefficient less than - 0.1 in the “Measured in” environment. One of these cells includes genes identified in the diauxic shift environment and measured in the cold stress environment (shown in more detail in panel E) and one includes a set of genes identified in the hydroxyurea shock environment and measured in the low phosphate environment (shown in more detail in panel F). (C-F) For each of the four cells highlighted with a yellow border and described above in panels A and B, the average expression of the gene set is shown for *S. cerevisiae* (yellow) and *S. paradoxus* (blue) in each of the six environments, with shading around each line indicating the 95% confidence interval of the mean. In each case, the red box surrounding a plot indicates the environment the gene set was identified in and the yellow box surrounding a plot indicates the environment this gene set was measured in that also showed a negative correlation less than -0.1 Note that these gene sets show highly correlated expression in all other environments they were measured in.

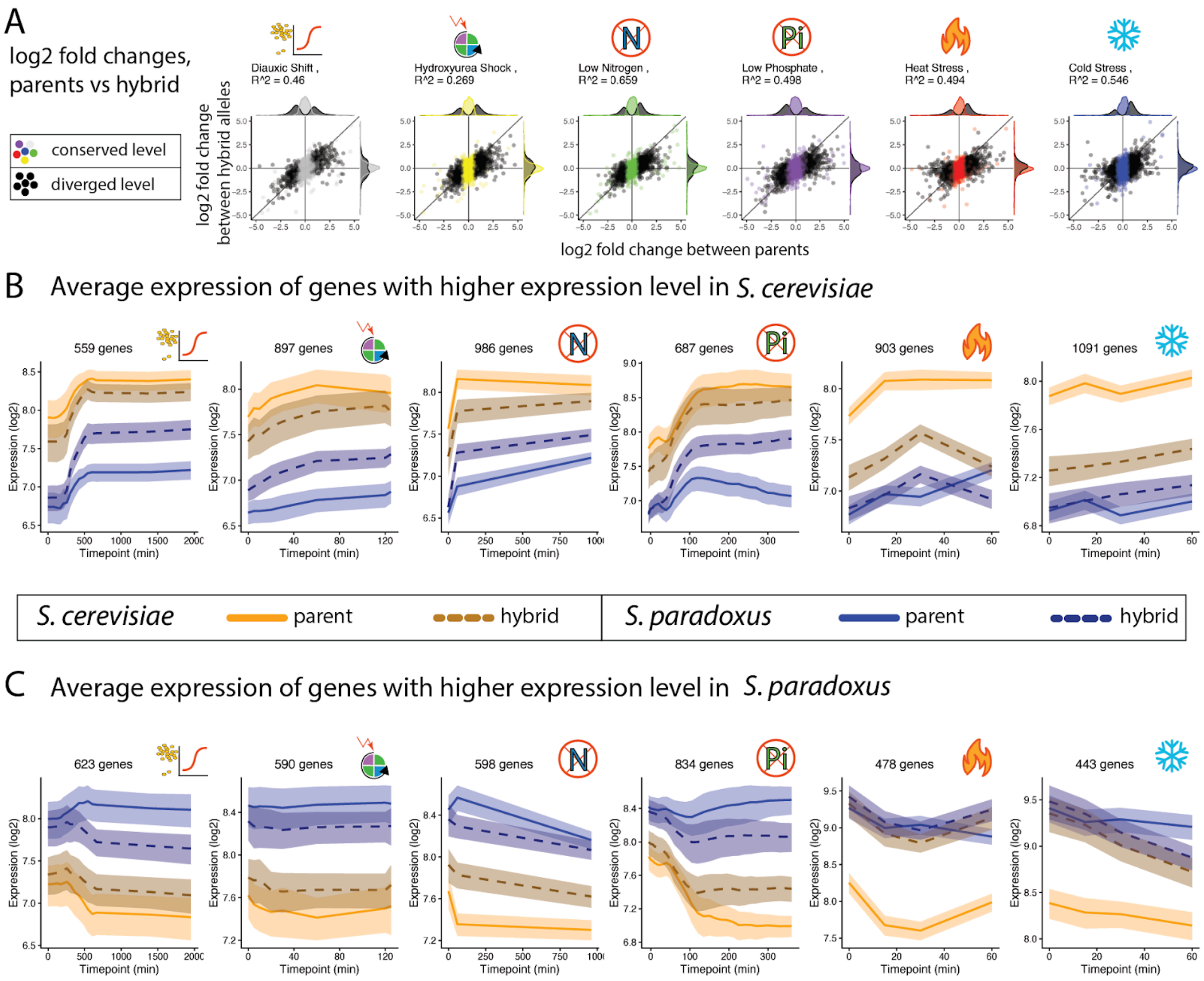


**Figure S6. Divergence in expression level is largely recapitulated in the $F_1$ hybrid allele-specific expression.** (A) Environment-specific plots comparing the $\log_2$ fold change in expression level between species to the $\log_2$ fold change in expression level between alleles in the $F_1$ hybrid. Logos representing the different environments are defined in Figure 2. These plots are similar to the plot in Figure 8B that shows these data combined for all six environments. The strong correlation between differences in expression level between species and between alleles in the $F_1$ hybrids suggests that *cis*-regulatory divergence is primarily responsible for divergence in expression level in all environments. (B) Average expression in *S. cerevisiae* (yellow) and *S. paradoxus* (blue), as well as expression of species-specific alleles in $F_1$ hybrids (dotted lines), is shown for genes with divergent expression levels that have higher expression in *S. cerevisiae*. Shading around each line indicates the 95% confidence interval of the mean. (C) Same as in (B) but for genes with divergent expression levels that have higher expression in *S. paradoxus*. Note that expression of species-specific alleles in the $F_1$ hybrid that are similar to the expression level of the parental species suggests *cis*-regulatory changes are primarily responsible for the divergence in expression level.

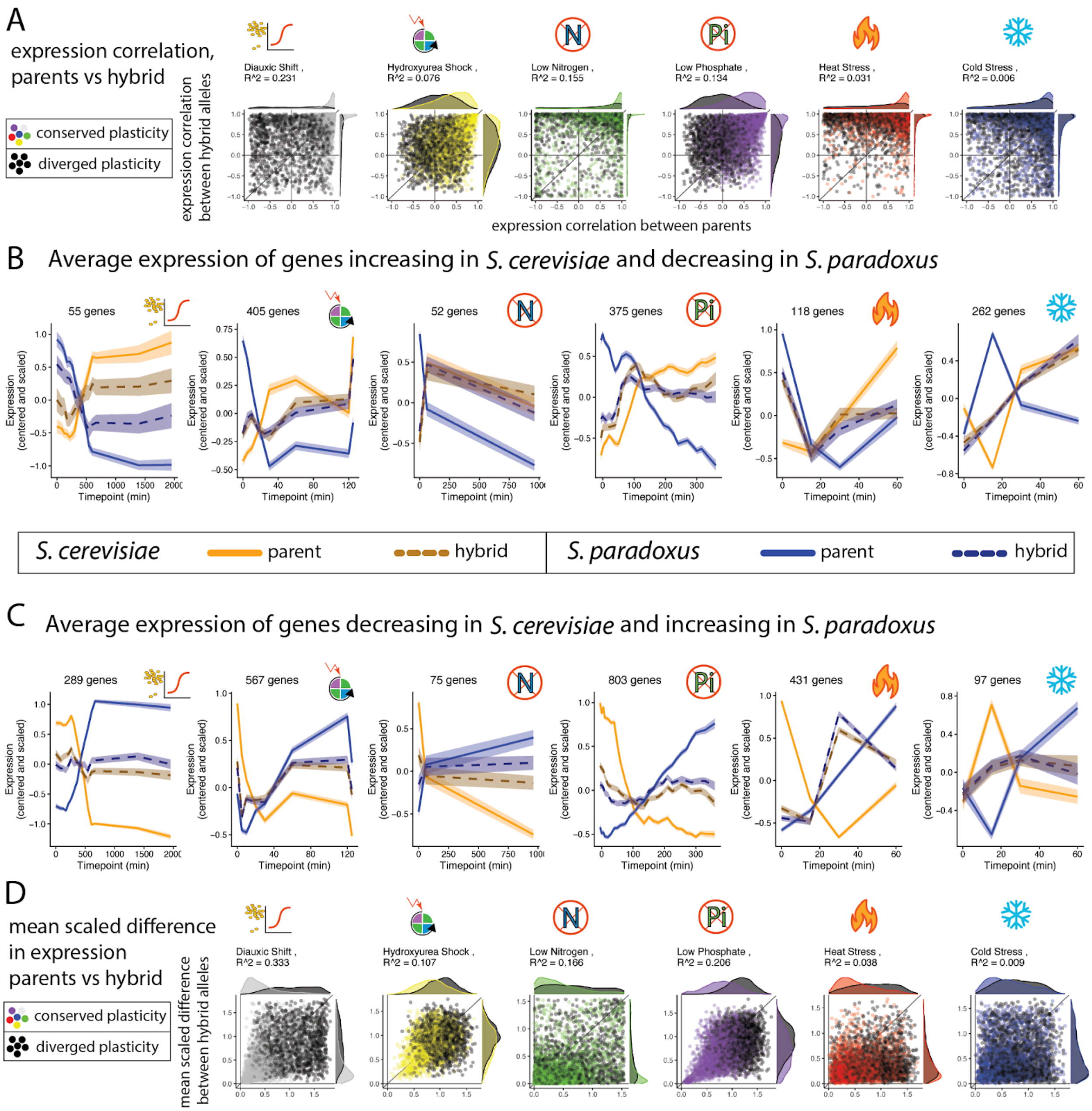


**Figure S7. Divergence in expression plasticity is rarely recapitulated in the $F_1$ hybrid allele-specific expression.** (A) Each scatterplot compares the expression correlation between parental species (*S. cerevisiae* and *S. paradoxus*) to the expression correlation between species-specific alleles in $F_1$ hybrids in a single environment. Logos representing the different environments are defined in Figure 2. These plots are similar to the plot in Figure 8C that shows these data combined for all six environments. The lack of a strong correlation between differences in expression plasticity between species and between alleles in the $F_1$ hybrids suggests that *trans*-regulatory divergence is primarily responsible for divergence in expression plasticity in all environments. (B) Average expression in *S. cerevisiae* (yellow) and *S. paradoxus* (blue), as well as expression of species-specific alleles in $F_1$ hybrids (dotted lines), is shown for genes with reversals in expression plasticity between species and expression increasing among timepoints within an environment in *S. cerevisiae*. Shading around each line indicates the 95% confidence interval of the mean. (C) Same as in (B) but for genes with reversals in expression plasticity between species that have expression decreasing across timepoints within an environment in *S. cerevisiae*. Note that expression of species-specific alleles in the $F_1$ hybrid that is more similar to each other than to expression in their parental species suggests *trans*-regulatory changes are primarily responsible for the divergence in expression plasticity. (D) These environment-specific plots are similar to (A) but use the mean scaled difference to compare expression between species and between alleles in $F_1$ hybrids rather than the correlation coefficients.

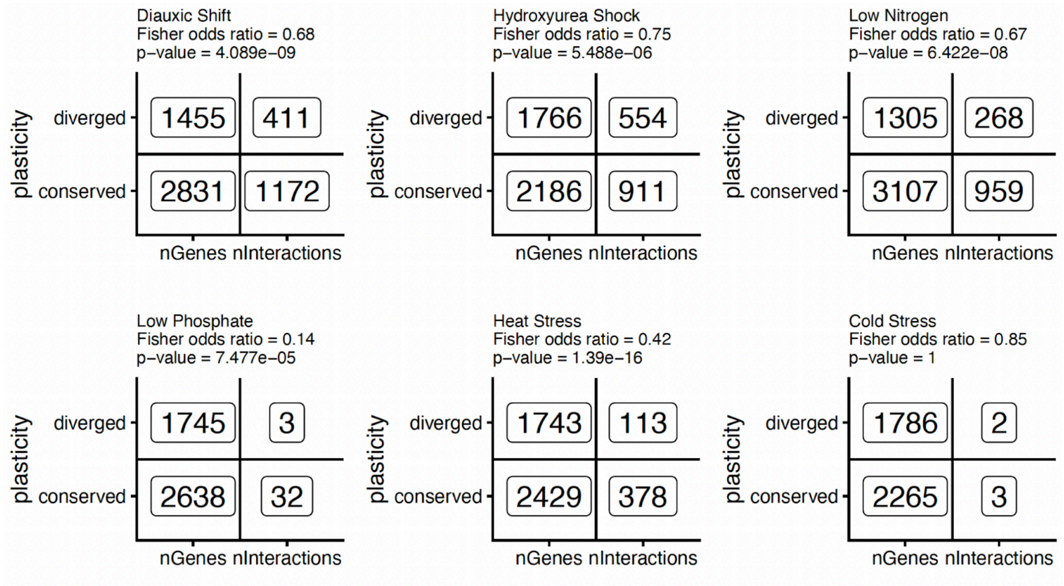


**Figure S8. Genes with conserved plasticity tend to have more regulatory connections in environment-specific networks.** Each 2x2 plot represents a Fisher Exact Test for enrichment in each environment. Genes were divided based on whether they had conserved or diverged in plasticity between *S. cerevisiae* and *S. paradoxus*. Regulatory interactions were divided by whether they targeted a gene with conserved or diverged plasticity.

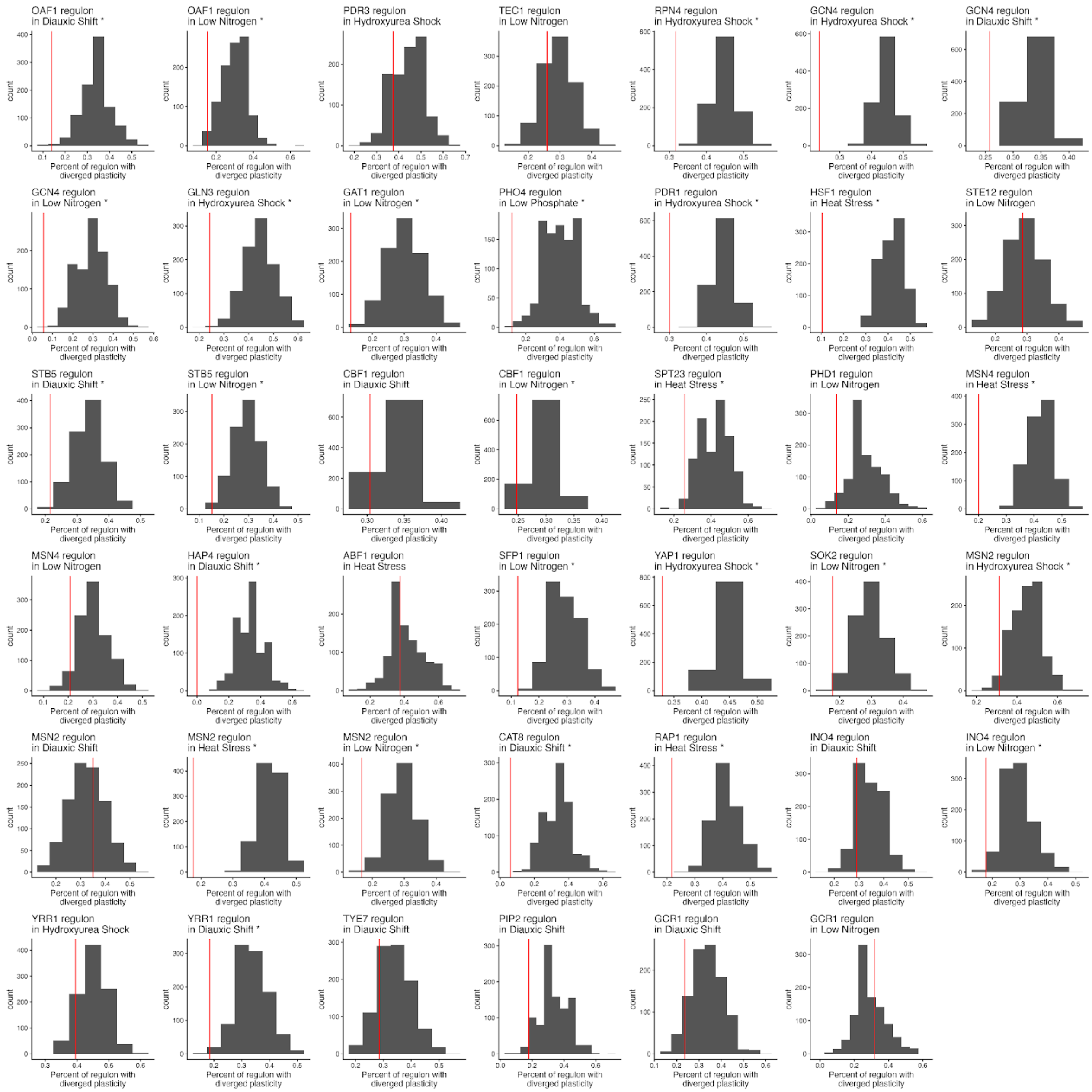


**Figure S9. Conservation of expression plasticity in environment-specific regulons with at least 20 genes.** Each plot shows the proportion of target genes with divergent plasticity for one regulator with divergent plasticity in one environment (vertical red line). The 41 plots shown correspond to the 41 regulators with at least 20 target genes in a specific environment, with the environment indicated. The grey histogram in each plot shows a null distribution generated by randomly sampling, 1000 times, the same number of genes as the number of target genes in that regulator's regulon. For each random sample, the percentage of genes with divergent plasticity in that environment was recorded and used to form the null distribution shown. Plots with asterisks after the name of the environment are regulons with less divergence in expression plasticity than at least 95% of the values in the null distribution.

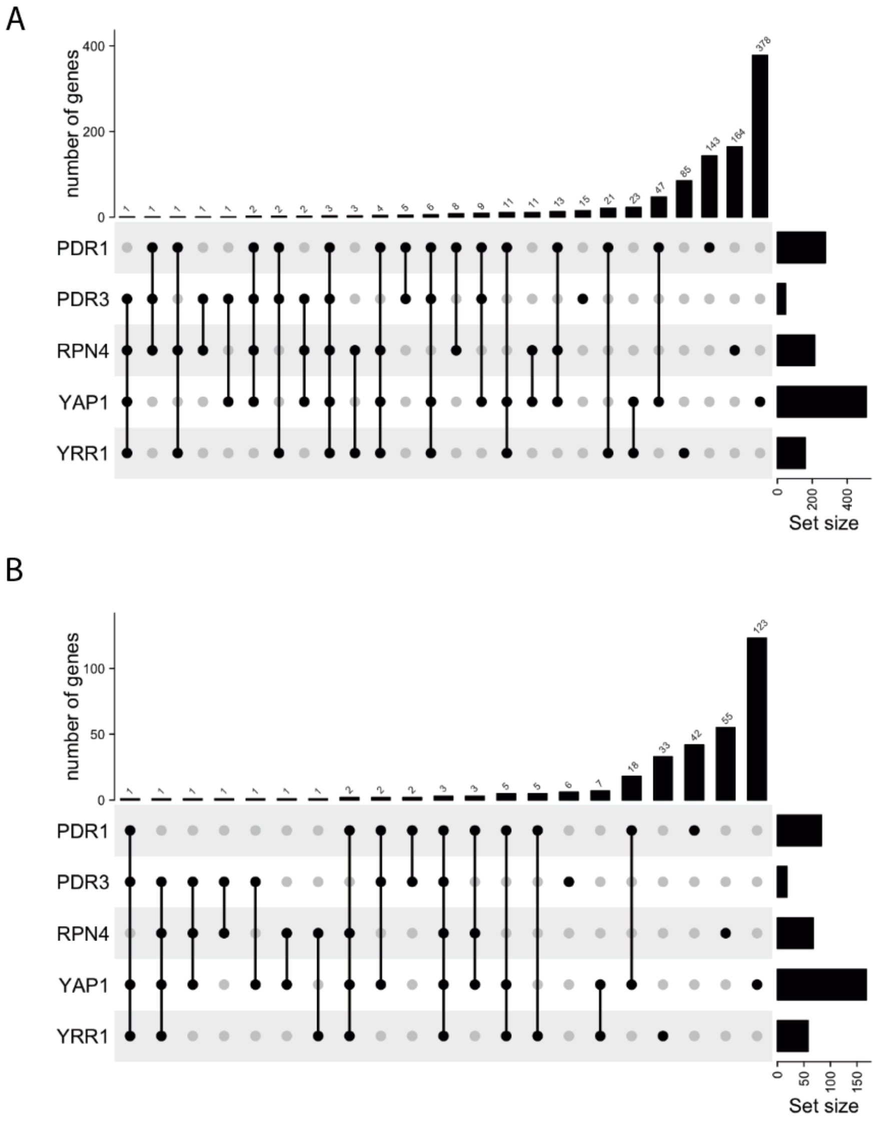


**Figure S10. Minimal overlap of genes with divergent expression plasticity that are targets of five transcription factors involved in the pleiotropic drug response network.** Considering all target genes of the transcription factors PDR1, PDR3, RPN4, YAP1, and YRR1 with evidence of divergent expression plasticity (among all six environments), the UpSet plot shows the number of shared target genes with divergent expression plasticity. The Set size shown to the right of the plot indicates the total number of genes with divergent expression plasticity for each individual transcription factor. The block dots connected by black lines show the sets of transcription factors considered for each of the bars in the bar plot. The number above each bar in the bar plot shows the number of genes with divergent expression plasticity that are target genes of the set of transcription factors indicated with black dots below the bar. A) All genes that were identified by Yeastract as a target of at least one of the five transcription factors. B) The subset of genes from panel A that we identified as having divergent plasticity in the environment with a documented regulatory interaction.
